# Vgll4 Proteins limit Organ Size in Zebrafish through Yap1-Dependent and -Independent Mechanisms

**DOI:** 10.1101/2025.05.08.652796

**Authors:** Alicia Lardennois, Chaitanya Dingare, Veronika Duda, Petra A Klemmt, Constanze Heinzen, David Kleinhans, Thorsten Falk, Carsten Schelmbauer, Olivia Mozolewska, Sofia Papadopoulou, Jason J K Lai, Didier Y R Stainier, Virginie Lecaudey

## Abstract

Precise control of organ size is crucial during development and homeostasis. Dysregulation of the underlying mechanisms can result in organ malformation and tumorigenesis. Although the Hippo signaling pathway plays a key role in regulating organ growth, the precise regulation of its effectors, YAP1 and WWTR1, remains unclear. To gain insights into tissue growth control during organ formation, we used the zebrafish posterior lateral line primordium (pLLP), a migratory group of epithelial cells that forms sensory organs, as a model. We demonstrate that Yap1 growth-promoting activity in the pLL system is modulated not only in the cytoplasm but also in the nucleus by Vgll4 proteins. We propose a model in which Yap1, together with Tead proteins, ensures that the pLLP contains a sufficient number of cells before migration begins. Vgll4b and Vgll4l, in contrast, function partially redundantly, to limit pLLP cell number, with Vgll4b showing a stronger tumor-suppressor activity. Our data indicate that Vgll4b/4l counteract Yap1 activity by competing with Yap1 for binding to Tead proteins, but also via a Yap1-independent mechanism. Altogether, this study reveals that a precise balance between Yap1 and Vgll4 proteins ensures proper regulation of cell number in the pLLP.

## Introduction

During embryonic development, it is crucial for organs to grow and acquire the correct size and shape. While cell proliferation is essential for organ formation, its dysregulation can lead to excessive growth and tumor formation. Therefore, proliferation must be precisely controlled and coordinated with other developmental processes to ensure that organs reach their proper size without exceeding it. Yet, the precise mechanisms underlying this regulation remain only partially understood.

The Hippo signaling pathway has emerged as a key regulator of organ size during development and homeostasis. Initially identified in *Drosophila* ^1–4^, the Hippo pathway is highly conserved in vertebrates ^5–7^. The canonical Hippo pathway is composed of a kinase cascade, which phosphorylates the downstream effectors YAP1 (Yes-associated protein 1) and its homolog WWTR1 (WW domain containing transcription regulator 1, also known as TAZ), leading to their sequestration in the cytoplasm or degradation. Conversely, when Hippo signaling is inactive, YAP1/WWTR1 translocate to the nucleus, where they drive the transcription of genes that promote proliferation and suppress apoptosis ^8–13^. The non-canonical pathway includes various upstream regulators, often membrane-associated cell junction proteins, that influence YAP1/ WWTR1 localization and stability, independently of their phosphorylation, at least in part. There is also substantial cross-talk between canonical and non-canonical pathways ^14^.

YAP1 and WWTR1 are transcription cofactors that lack a DNA-binding domain. They need to interact with DNA-binding proteins, particularly TEAD-family transcription factors, to regulate the expression of target genes. TEAD proteins also interact with Vestigial-like family member 4 (VGLL4), which functions as a tumor suppressor ^15–17^. Several studies have shown that VGLL4 and YAP1 compete for binding to TEAD, because they share the same binding region ^15, 18, 19^. The competition for TEAD binding has been proposed to determine the outcome of gene expression. Elevated levels of VGLL4 competitively displace YAP1 from TEAD, leading to the downregulation of genes associated with cell proliferation and survival, accounting for the tumor-suppressive role of VGLL4 ^20, 21^. Conversely, when YAP1 predominates over VGLL4 and binds to TEAD, it activates the expression of genes promoting cell proliferation and survival. In this scenario, the balance is shifted towards promoting cell growth and potentially contributing to tumor development ^15, 17^. Although VGLL4 tumor suppressor activity has mainly been associated with the capacity of VGLL4 to compete with YAP1 for binding to TEAD, it remains unclear whether this is the only function of VGLL4 or whether VGLL4 has YAP1-independent functions. Indeed, VGLL4 has also been proposed to function together with TEADs as a default repressor, the main function of YAP1 in this context being to counteract this repression ^19, 22^. Thus, the mechanisms that precisely regulate the Hippo pathway output are not yet fully understood.

The simplicity and robustness of the zebrafish posterior lateral line primordium (pLLP) provides an excellent model to investigate this question further. As it delaminates from the pLL placode, the pLLP is composed of approximately 120 epithelial cells that migrate as a group just underneath the skin and deposit neuromasts at their trailing end ^23, 24^. Its compact size, robust cell number at any given time and superficial position, just below the flat skin cells, make the pLLP an ideal system for high-resolution imaging, allowing one to automatically quantify the number of cells as well as various parameters such as their shape and volume. Combined to the amenability of zebrafish embryos to genetic and pharmacological approaches, the model has obvious advantages for dissecting the molecular mechanisms controlling epithelial tissue growth. The canonical Wnt signaling pathway, via Lef1, has been shown to promote proliferation in the leading region, thus partially compensating for the loss of cells due to neuromast deposition ^25–29^. We have previously reported that Yap1 is required for the pLLP to reach a sufficient number of cells, and that the Motin family protein Amotl2a is necessary to limit the number of cells in pLLP by inhibiting Yap1 and Lef1 activity ^30^. However, the mechanisms that regulate the precise number of cells in the migrating pLLP are not completely understood, in particular the mechanisms limiting the activity of Yap1 in the cytoplasm and in the nucleus.

In the current work, we investigated the mode of action of Yap1 in the nucleus and its regulation by Vgll4 using the zebrafish pLLP as an *in vivo* model. We demonstrate that Yap1 growth promoting activity in the pLL system is TEAD-dependent and is already required in the pLL placode, before the onset of pLLP migration. By combining loss-of-function, rescue and epistasis experiments with reporter lines and quantitative imaging, we demonstrate that *vgll4b* and *vgll4l*, the two zebrafish *vgll4* paralogs expressed in the pLLP, function partially redundantly to limit the size and cell number in the pLLP. Our data further indicate that the tumor-suppressor function of Vgll4 in the pLLP is also Tead-dependent, and that Vgll4b and Vgll4l limits pLLP cell number by competing with Yap1 to bind to Tead, but also in a Yap1-independent manner.

## Results

### 1. The growth-promoting activity of Yap1 in the pLLP depends on Tead proteins

To test whether the growth-promoting function of Yap1 in the pLLP depends on Tead transcription factors, we took advantage of a *yap1* allele, *yap1^bns^*^22^, encoding a Yap1 protein lacking 18 amino acids in the predicted Tead-binding domain (Fig. 1A) ^31^. The deletion includes the Serine at position 54 (S54) which is required for the interaction with Tead ^32–34^. We first quantified the size and shape of the pLLP in *yap1^bns^*^22^ mutant and compared them to those in *yap1^fu^*^47^ and *yap1^fu^*^48^, two other mutant alleles carrying a premature STOP codon that we previously generated (Fig. 1A) ^30^. Similar to *yap1^fu^*^47^ and *yap1^fu^*^48^, *yap1^bns^*^22^ mutant pLLP appeared smaller and rounder (Fig. 1B-E). Quantification confirmed a reduction in the volume of the pLLP (Supplementary Fig. 1A), as well as in the cell count of approximately 20% for all three alleles compared to their respective siblings (Fig. 1F). In addition to the reduction in size, mutant pLLP were rounder than their siblings for all three alleles, as shown by a significant reduction in the aspect ratio (Fig. 1G). The number of rosettes was reduced in all three alleles, whereas the ‘rosettiness’, which quantifies the shape and quality of the rosettes (see Materials and methods), was not reduced (Supplementary Fig. 1C-D). This suggests that primordia containing fewer cells form rosettes of similar size and shape, but that there are fewer of them.

**Figure 1.**
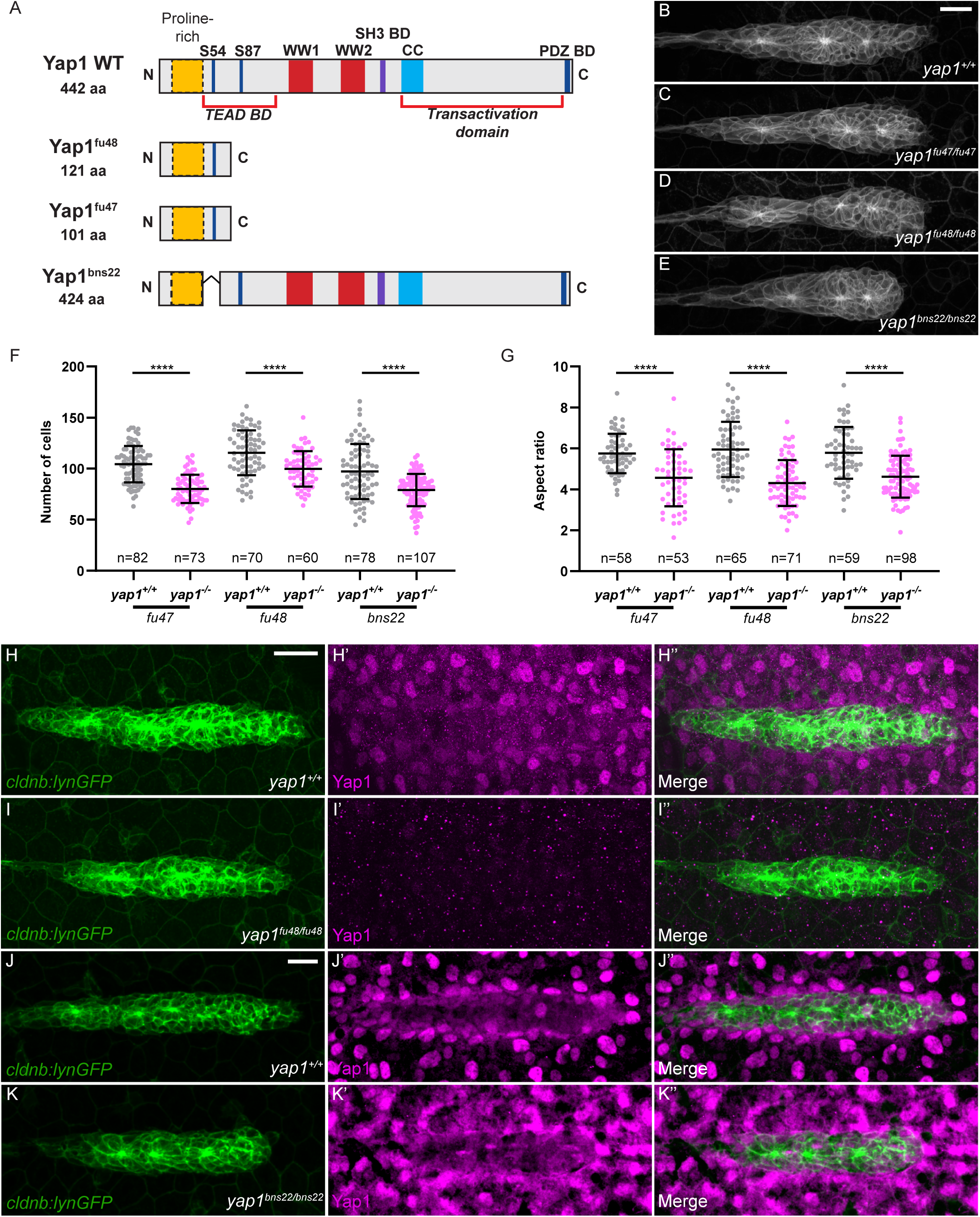
Tead-binding is essential for Yap1-mediated proliferation in the pLLP. (A) Schematic representation of the wild-type Yap1 and the mutant Yap1^fu47^, Yap1^fu48^ and Yap1^bns22^ proteins. (B-E) Maximum intensity projection (MIP) of spinning disc confocal Z-stacks showing the pLLP in wild-type (B), *yap1^fu47^* (C), *yap1^fu48^* (D) and *yap1^bns22^* (E) homozygous mutant embryos. (F-G) Quantification of cell number (F) and aspect ratio (G) of the pLLP across genotypes. (H-K’’) MIP of confocal Z-stacks showing Yap1 localization in control (H-H’’, J-J’’), *yap1^fu48^* (I-I’’) and *yap1^bns22^* mutant embryos (K-K’’). Unless otherwise indicated, the pLLP is marked by *cldnb:lynGFP* in all panels (B-K’’). Scale bars: 20 μm (B-E, H-K’’). In all figures MIP are maximum intensity projection of spinning disc confocal Z-stacks

To ensure that we were specifically testing the requirement for Tead binding, we confirmed that the mutant Yap1^bns^^22^ protein was effectively produced and stable. We first quantified the amount of *yap1* mRNA in each mutant allele by qRT-PCR. While *yap1^fu47^* and *yap1^fu48^* mutant mRNAs were significantly reduced (56 and 65%, respectively), *yap1^bns22^* mutant mRNA levels remained unchanged (Supplementary Fig. 1B). These results suggest that *yap1^fu47^* and *yap1^fu48^* mRNAs, but not *yap1^bns22^*, are degraded, most likely by nonsense-mediated mRNA decay (NMD) due to the presence of premature STOP codons ^35^. To measure the level of the corresponding proteins, we generated an antibody against the zebrafish Yap1 protein (VL7801) ^36, 37^. Immunofluorescence staining of wild-type (WT) embryos at 32 hours post-fertilization (hpf) showed that Yap1 was present at a low level and homogenously in the cells of the pLLP, whereas the neighboring somitic cells showed distinct nuclear localization (Fig. 1H-H’’). There was no signal in *yap1^fu^*^47^*^/fu47^*(not shown) nor in *yap1^fu48/fu48^* (Fig. 1I-I’’) mutants, confirming the specificity of the antibody and that these are null mutants. In contrast, Yap1 proteins were still present in *yap1^bns22/bns22^* mutants, at levels at least equivalent to WT Yap1 (Fig. 1J-K’’). The signal appeared more diffuse in *yap1^bns22/bns22^* mutants, suggesting a shift of the Yap1^bns22^ mutant protein towards the cytoplasm both in the pLLP and in the surrounding somites (Fig. 1J’,K’). This is in agreement with published data showing that binding to TEAD retains YAP1 in the nucleus ^38^. Taken together, these results indicate that Yap1-Tead interaction is essential for the pLLP to have an appropriate cell number and shape, but not for rosette assembly.

To further confirm that Yap1-Tead interaction was important for the pLLP to reach its ‘normal’ size, we undertook a classical genetic approach to rescue *yap1* mutant phenotype with the WT form of Yap1 and a Tead-binding defective form of Yap1 in which the Serine in position 54 is mutated in Alanine (*yap1^S54A^*) ^32^. While *yap1^WT^*mRNA fully rescued both cell number and aspect ratio to control levels (Fig. 2A,C-D,G-H), *yap1^S54A^*mRNA did not rescue the reduced cell number or the reduced aspect ratio in *MZyap1^fu48/fu48^*embryos (Fig. 2A,E-F,I-J). Neither *yap1^WT^* nor *yap1^S54A^*overexpression in WT induced any obvious phenotype or altered cell numbers in the pLLP (Fig. 2A-B,E,G,I). Similar results were obtained when rescuing the *yap^bns22^* mutant (Supplementary Fig. 2A-F). Altogether, these results show that Yap1 needs to physically interact with Tead to maintain a proper number of cells in the pLLP.

**Figure 2.**
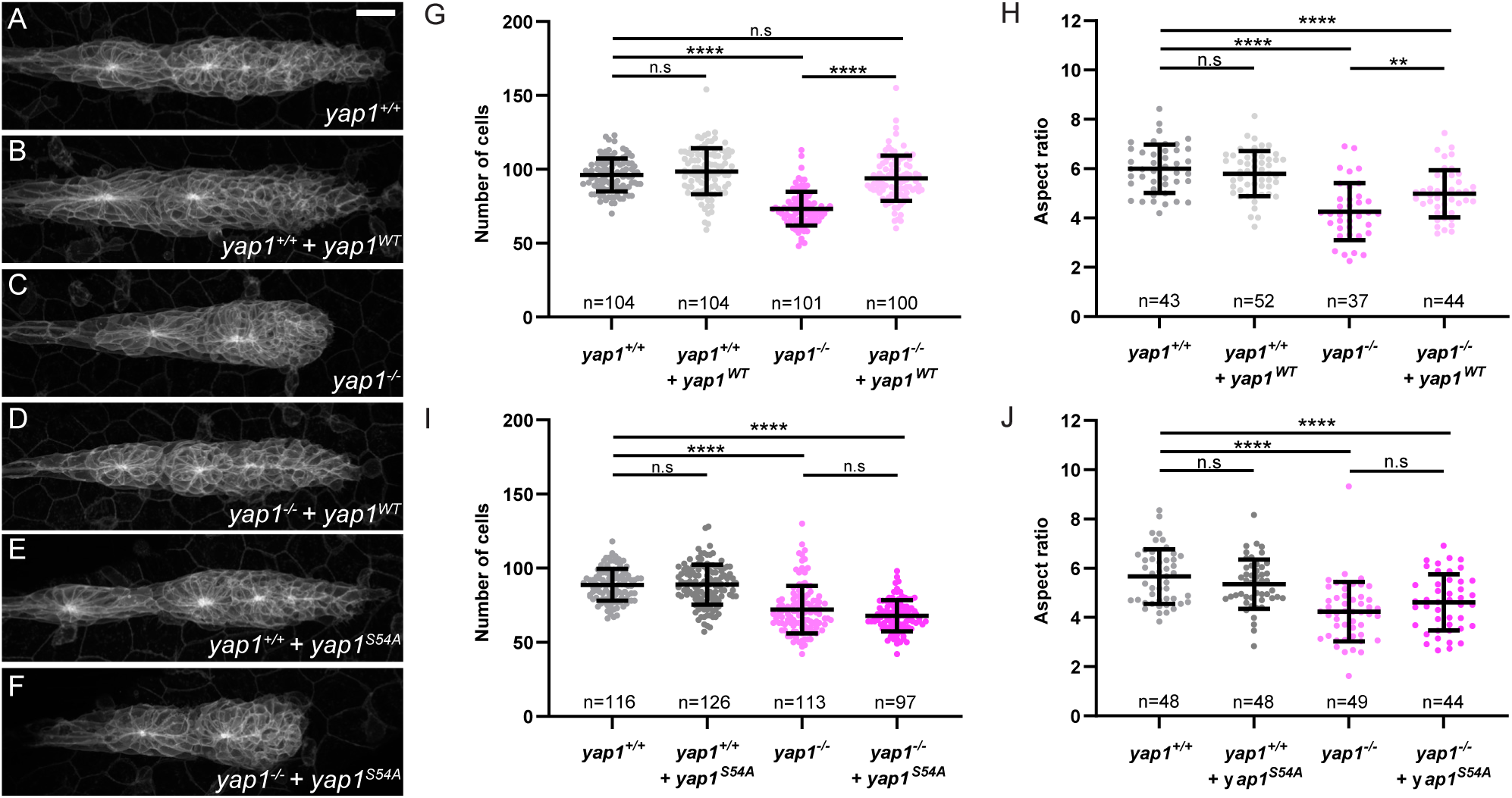
A Tead-Binding-Deficient Yap1 mutant fails to rescue *yap1* loss-of-function phenotype. (A-F) MIP of confocal Z-stacks showing differences in pLLP size in uninjected *yap1^+/+^* embryos (A), *yap1^+/+^* embryos injected with *yap1*-WT mRNA (B), *yap1^+/+^* embryos injected with *yap1-S54A* mRNA (C), uninjected *yap1^-/-^* embryos (D), *yap1^-/-^* embryos injected with *yap1*-WT mRNA (E), *yap1^- /-^* embryos injected with *yap1-S54A* mRNA (F). (G-J) Quantification of pLLP cell number (G, I) and aspect ratio (H, J) of the pLLP in the indicated conditions. Scale bar: 20 μm.

### 2. Vgll4b and Vgll4l are required to limit the number of cells in the pLLP

Vestigial-like family member 4 (VGLL4) has been identified, both in *Drosophila* (Tgi) and in mammals, as a transcriptional cofactor that competes with YAP1 for binding to TEAD transcription factors ^19, 22^. Thus, we wanted to determine whether Vgll4 proteins similarly compete with Yap1 activity in the pLLP. We first assessed the expression patterns of the three zebrafish *vgll4* paralogs in the embryos, in particular *vgll4b* and *vgll4l*, which have been reported to be expressed in pLL neuromasts ^39^. *vgll4b* and *vgll4l* were expressed in the pLLP at 30 hpf (Fig. 3A-D’) and until the end of migration (not shown). In contrast, *vgll4a* was not expressed in the pLLP (Supplementary Fig. 3C). We then tested whether the two *vgll4* paralogs expressed in the pLLP might be involved in controlling its cell count and size. For this, we generated mutants for *vgll4b* and *vgll4l* using TALEN- and CRISPR-mediated mutagenesis, respectively. The recovered alleles carried deletions of 13 bp and 11 bp, respectively, leading to premature STOP codons in the first quarter of the proteins in both cases (Fig. 3E-F, Supplementary Fig. 3A-B). *vgll4b* and *vgll4l* homozygous mutant embryos did not display any obvious morphological defects and the adults were viable. This is in agreement with other recently published mutants of these genes ^40, 41^. Next, we quantified the number of cells in *vgll4b* and *vgll4l* mutant pLLP. While *vgll4l* embryos did not show any phenotype on the pLLP cell count, the *vgll4b* pLLP had significantly more cells with a mean increase of 35% as compared to control siblings (Fig. 3G-I,M). To test whether NMD-mediated transcriptional adaptation may limit or mask the phenotype of *vgll4b* and *vgll4l* mutants, we quantified the amount of *vgll4b* and *vgll4l* mRNA in both mutants. *vgll4b* mRNA was reduced by approximately 60% in *vgll4b* mutant, possibly because of NMD, but *vgll4l* mRNA levels were not increased (Supplementary Fig. 3G). In contrast, *vgll4l* mRNA was increased two-fold in *vgll4l* mutants, suggesting that Vgll4l directly or indirectly inhibits its own expression and is not degraded. *vgll4b* mRNA levels were also not increased in *vgll4l* mutants (Supplementary Fig. 3H). This indicates that the loss of Vgll4b or Vgll4l activity is not compensated by an upregulation of *vgll4l* or *vgll4b* mRNA, respectively.

**Figure 3.**
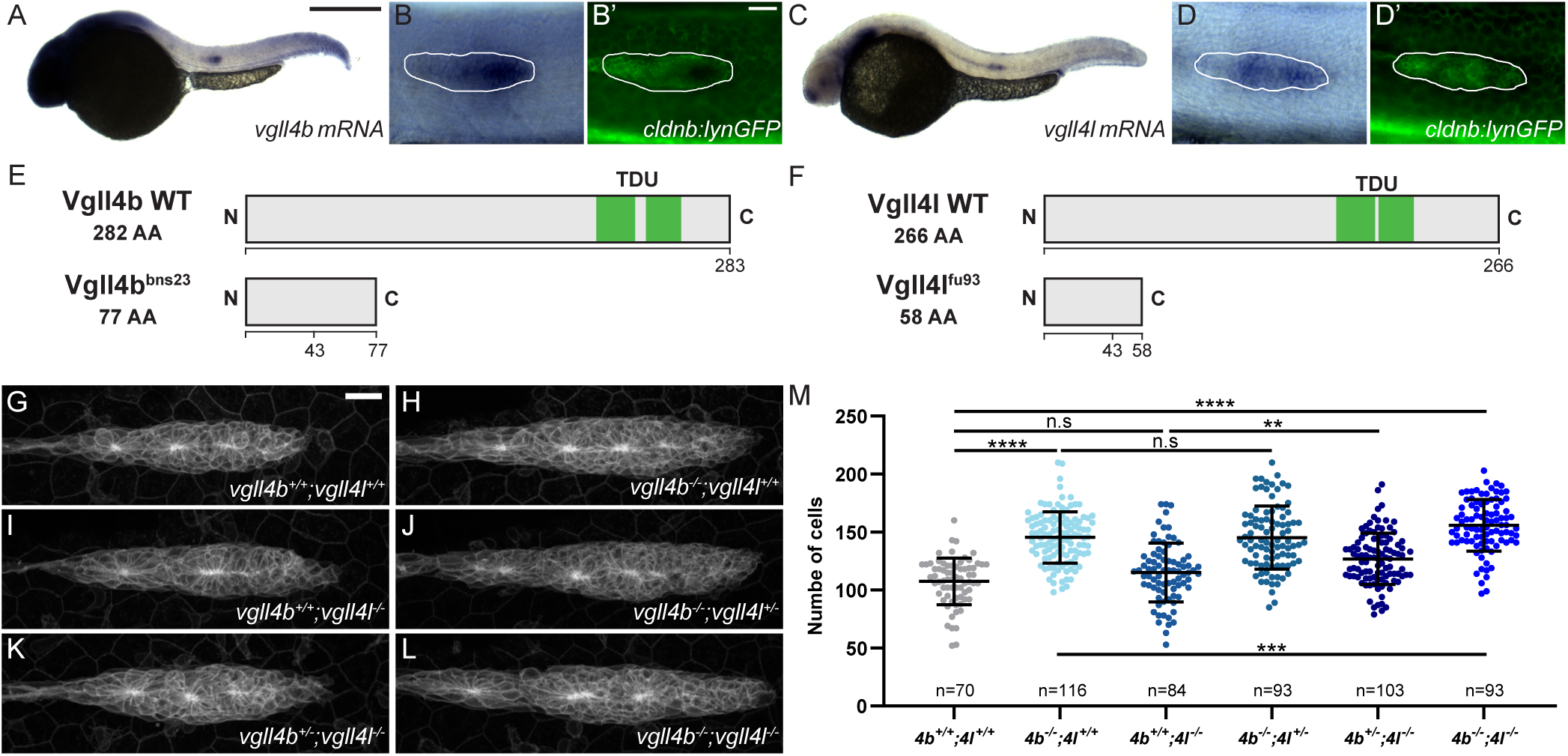
Vgll4b and Vgll4l are required to limit the number of cells in the pLLP. (A-D’) Overview (A, C) and higher magnification images (B, B’, D, D’) taken on a brightfield microscope of 32hpf *cldnb:lynGFP* embryos stained by in situ hybridization (ISH) for *vgll4b* (A-B’) and *vgll4l* (C-D’) and counterstained with an anti-GFP antibody (B’, D’). (E-F) Schematic representation of wild-type Vgll4b and the mutant form Vgll4b^bns23^ (E), and wild-type Vgll4l and the mutant form Vgll4l^fu93^ (F). (G-L) MIP of confocal Z-stacks showing the pLLP in *vgll4b^+/+^;vgll4l^+/+^*(G), *vgll4b^-/-^;vgll4l^+/+^* (H), *vgll4b^+/+^;vgll4l^-/-^* (I), *vgll4b^-/-^;vgll4l^+/-^* (J), *vgll4b^+/-^;vgll4l^-/-^* (K), *vgll4b^-/-^;vgll4l^-/-^* (L) embryos. (M) Quantifications of cell number in the pLLP in each indicated group. Scale bar: 400 μm (A, C), 20 μm (B-D’, G-L).

Since both genes are expressed in the pLLP, they could still functionally compensate for each other’s loss. To test this, we generated *vgll4b*;*vgll4l* double mutants. The number of cells in the *vgll4b*;*vgll4l* pLLP was not only higher than in control siblings (mean increase of 50%, p<0.0001), but was also significantly higher than in *vgll4b* single mutants (Fig. 3G-M). Quantification of the pLLP volumes showed the same trend as the cell counts in both single and double mutants (Supplementary Fig. 3I), indicating that the increase in cell counts translates to an increase in organ size. No difference in the aspect ratio or in the number and quality of the rosette could be detected in any of the *vgll4* mutants (Supplementary Fig. 3J-L). The final pattern of neuromasts was not significantly affected, despite the increase in cell number in the pLLP (Supplementary Fig. 3M-R). To exclude the possibility that *vgll4a* may compensate for vgll4b/4l loss-of-function, we confirmed that *vgll4a* was not expressed in the double mutants using in-situ hybridization (ISH) (Supplementary Fig. 3C-F’’). Taken together, these results show that Vgll4l and Vgll4b function largely redundantly and are required to limit the number of cells in the pLLP.

### 3. Vgll4b is sufficient to reduce the number of cells in the pLLP and to rescue the absence of Vgll4 activity in the pLLP

To gain deeper insight into the function of Vgll4b in limiting pLLP cell number, we performed overexpression and rescue experiments using WT Vgll4l and Vgll4b mRNA. As expected, the tRFP-Vgll4b fusion protein was localized to the nucleus, including in pLLP cells (Supplementary Fig. 4C’). Overexpression of *vgll4b* mRNA in WT embryos resulted in an approximately 20% reduction in pLLP cell number, indicating that Vgll4b is not only required but also sufficient to limit cell numbers in the pLLP (Fig. 4A,B,G). Furthermore, injection of WT *vgll4b* mRNA in *vgll4b^-/-^* single mutants or *vgll4b^-/-^*;*vgll4l^-/-^* double mutant embryos fully rescued the increase in pLLP cell number to control levels (Fig. 4A,C-G). Quantification of the total primordium volume confirmed these observations (Supplementary Fig. 4A). Taken together, these results indicate that Vgll4b is necessary and sufficient to limit the number of cells in the pLLP and that it is also sufficient to rescue the entire loss of Vgll4 activity in the pLLP.

**Figure 4.**
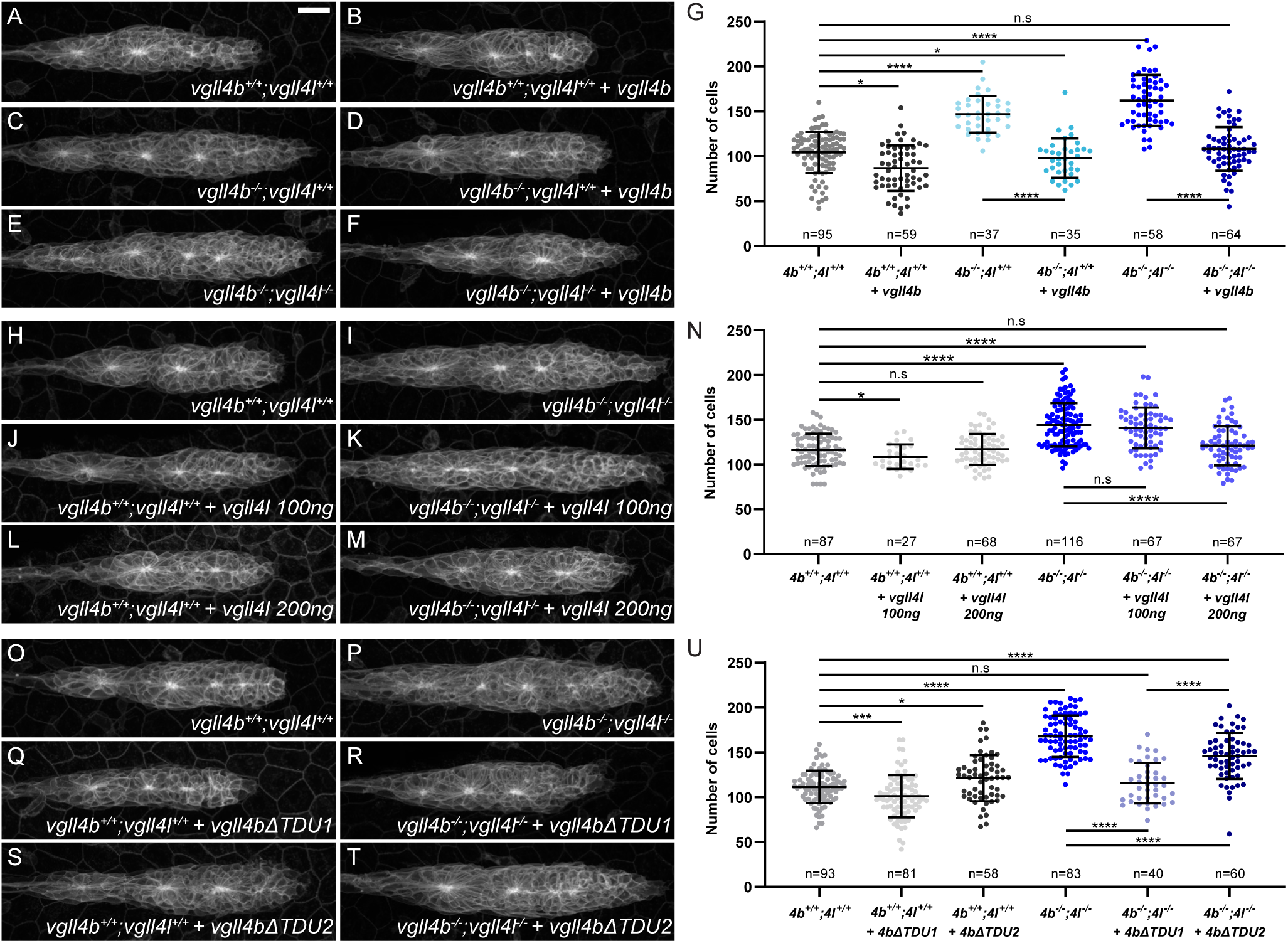
Vgll4b and Vgll4l are sufficient to rescue the loss of Vgll4 activity in the pLLP, and Vgll4b TDU2 is required for this function. (A-F) MIP of confocal Z-stacks showing the pLLP in *vgll4b^+/+^;vgll4l^+/+^* (A), *vgll4b^-/-^;vgll4l^+/+^* (C), *vgll4b^-/-^;vgll4l^-/-^* (E) embryos and the same genotypes injected with *tRFP-vgll4b* mRNA (B, D, F). (G) Quantification of pLLP cell number in each condition. (H-M) MIP of confocal Z-stacks showing the pLLP in *vgll4b^+/+^;vgll4l^+/+^*(H), and *vgll4b^-/-^;vgll4l^-/-^* (I) embryos, injected with *vgll4l* mRNA at 100ng/ μL (J, K) or at 200ng/μL (L, M). (N) Quantification of pLLP cell number in each condition. (O-T) MIP of confocal Z-stacks showing the pLLP in *vgll4b^+/+^;vgll4l^+/+^* (O), and *vgll4b^-/-^;vgll4l^-/-^* (P) embryos, injected with *tRFP-vgll4bΔTDU1* (Q, R) or *tRFP-vgll4bΔTDU2* (S, T) mRNA. (U) Quantification of pLLP cell number in each condition. Scale bar: 20 μm.

In contrast, injection of *vgll4l* mRNA at the same effective concentration as *vgll4b* had no effect on the pLLP cell number in WT embryos (Fig. 4H,J,N) and failed to rescue the increased cell count in *vgll4b^-/-^; vgll4l^-/-^* mutants (Fig. 4I,K,N). Doubling the concentration of *vgll4l* mRNA fully rescued the double mutant phenotype (Fig. 4I,M,N) without affecting the pLLP cell number in WT embryos (Fig. 4H,L,N). These results indicate that while both Vgll4b and Vgll4l can restore Vgll4 function in the pLLP, Vgll4b exhibits greater potency. Consistent with this, Vgll4b, but not Vgll4l, was sufficient to restrict pLLP cell number.

### 4. The TDU domains are differentially required for the tumor suppressor activity of Vgll4b

VGLL4 is the only member of the VGLL family containing two Tondu (TDU) motifs. The TDU motifs, in particular TDU2, have been shown to be necessary and, at least in some contexts, sufficient for the interaction of VGLL4 with TEAD, and for VGLL4 growth-inhibitory function ^15, 17, 21^. To test whether the growth-limiting activity of Vgll4b in the pLLP depends on its interaction with Tead proteins, we generated *vgll4b* constructs lacking either the first or the second TDU domain (Supplementary Fig. 4F, G). Both variants localized to the nucleus, similar to WT Vgll4b, indicating that neither TDU motif is essential for Vgll4b nuclear localization (Supplementary Fig. 4C-E’). Injecting *vgll4bΔTDU1* mRNA in WT resulted in a similar reduction in cell number as compared to the overexpression of WT *vgll4b* (approximately 10%) and to a full rescue of the phenotype of the *vgll4b^-/-^*;*vgll4l^-/-^* double mutants (Fig. 4O-R, U). In contrast, injecting *vgll4bΔTDU2* in WT embryos did not lead to a decreased number of cells, but even to a slight increase (approximately 9%) and resulted only in a partial rescue of the *vgll4b^-/-^*;*vgll4l^-/-^* double mutant phenotype (Fig. 4O-P, S-U). These results indicate that, similar to Yap1, the growth-limiting activity of Vgll4b in the pLLP is Tead-dependent and that the Vgll4b-Tead interaction is largely mediated by TDU2.

### 5. Competitive and independent functions of Vgll4 and Yap1 in Tead binding and regulation of cell proliferation Vgll4 and Yap1 compete for binding to Tead

So far, our results show that the activity of Vgll4b in the primordium depends on its interaction with Tead and therefore probably relies on a competitive interaction with Yap1 to bind to Tead as shown in other contexts ^15, 19, 22^. If this is the case, Yap1 activity should increase in embryos lacking Vgll4 activity. To test this hypothesis, we overexpressed *yap1* in the *vgll4* mutants. As previously shown (Fig. 1F), overexpressing *yap1* in control *vgll4b^+/+^;vgll4l^+/+^*embryos was not sufficient to increase pLLP cell number (Fig. 5A-B, I). In contrast, injection of *yap1* mRNA in *vgll4b^-/-^* single or in *vgll4b^-/-^*;*vgll4l^-/-^* double mutants lead, respectively, to a 20% and 25% increase in pLLP cell counts compared to the respective non-injected mutants (Fig. 5C-D, G-H, I). Overexpression of *yap1* in *vgll4l^-/-^* single mutants did not lead to an increase in the cell number (Fig. 5E-F, I). Altogether, these results indicate that Yap1 activity is enhanced in the absence of Vgll4, supporting a competitive interaction between Vgll4 and Yap1 in the migrating pLLP. They also confirm that the functional redundancy between *vgll4b* and *vgll4l* is not symmetrical: while the loss of Vgll4l activity alone is not sufficient to make Yap1 more potent, the absence of Vgll4b alone is.

**Figure 5.**
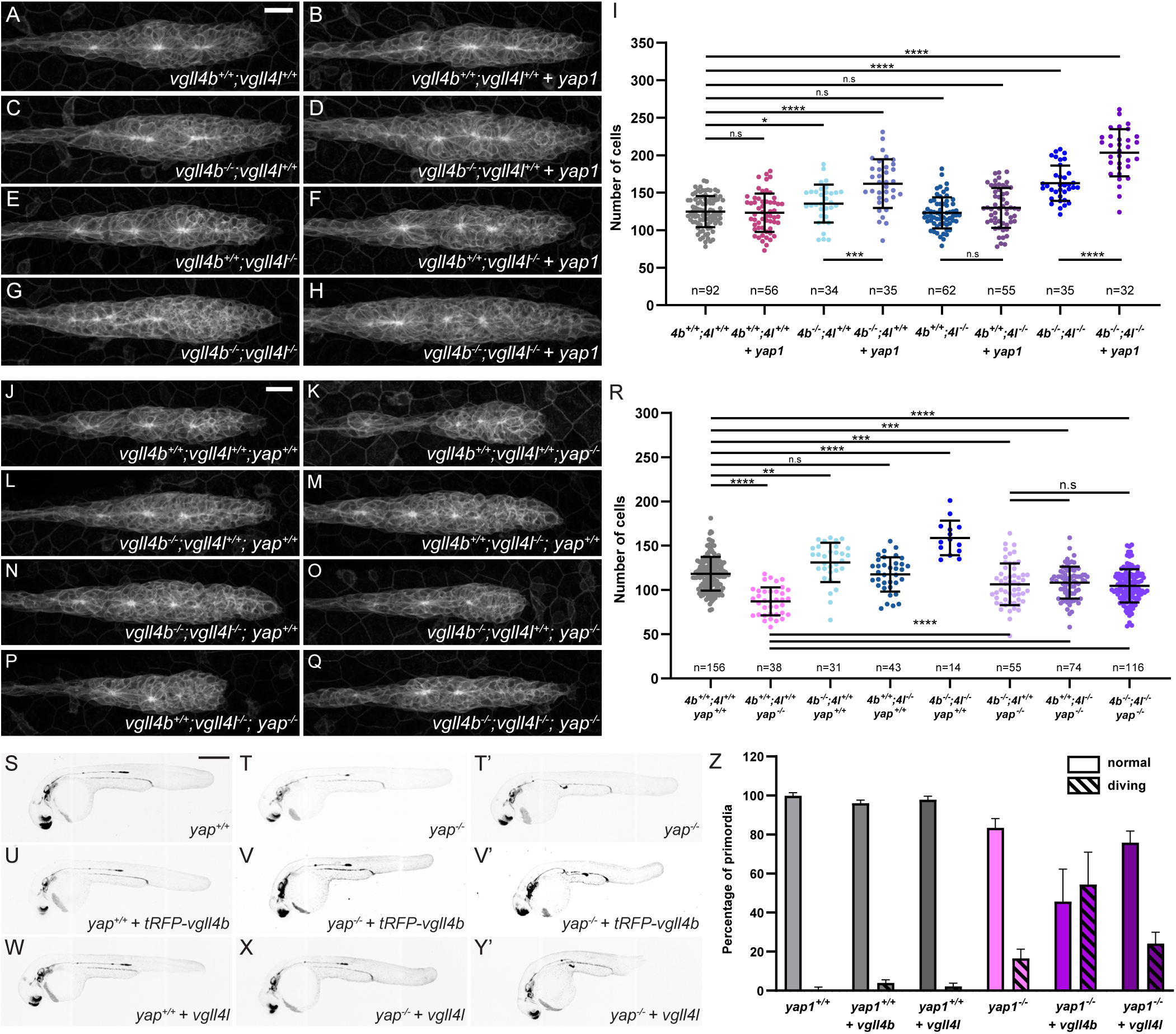
Vgll4 limits cell number in the pLLP by competing with Yap1 for binding to Tead, but also through a Yap1-independent mechanism. (A-H) MIP of confocal Z-stacks showing the pLLP in *vgll4b^+/+^;vgll4l^+/+^*(A), *vgll4b^-/^;^-^vgll4l^+/+^* (C), *vgll4b^+/+^;vgll4l^-/-^*(E), *vgll4b^-/-^;vgll4l^-/-^* (G) embryos and the same genotypes, respectively, injected with *tBFP-yap1* mRNA (B, D, F, H). (I) Quantification of pLLP cell number in each condition. (J-Q) MIP of confocal Z-stacks showing the pLLP in *vgll4b^+/+^;vgll4l^+/+^;yap1^+/+^* (J), *vgll4b^+/+^;vgll4l^+/+^;yap1*^- /-^ (K), *vgll4b^-/-^;vgll4l^+/+^;yap1^+/+^* (L), *vgll4b^+/+^;vgll4l^-/-^;yap1^+/+^* (M); *vgll4b^-/-^;vgll4l^-/-^;yap1^+/+^* (N), *vgll4b^- /-^;vgll4l^+/+^;yap1^-/-^*(O), *vgll4b^+/+^;vgll4l^+/+^;yap1^-/-^* (P), *vgll4b^-/-^;vgll4l^-/-^;yap1^-/-^* (Q) embryos. (R) Quantification of pLLP cell number in each condition. (S-Y’) MIP of confocal Z-stacks showing overview images of 32hpf *yap1^+/+^* (S), *yap1^-/-^* (T, T’), and embryos of the same genotypes injected with *tBFP-vgll4b* mRNA (U-V’) or with *vgll4l* mRNA (W-Y’). (Z) Percentage of pLLP displaying normal migration or “pLLP diving” in each condition. Scale bars: 20 μm (A-H, J-Q) and 400 μm (S- Y’).

#### Vgll4b and Vgll4l exhibit a function independent of Yap1

To further investigate the competition between Vgll4 and Yap1, we generated *vgll4b;yap1, vgll4l;yap1* double and *vgll4b;vgll4l;yap1* triple mutants. We reasoned that if Vgll4b and Vgll4l function exclusively by competing with Yap1 to bind to Tead, a combined loss of Yap1 and Vgll4 function should result in a phenotype similar to *yap1* single mutants. Essentially, in the absence of Yap1, Vgll4 would not be able to complete its function of Yap1 competitor. Notably, *vgll4b^-/-^;yap1^-/-^, vgll4l^-/-^;yap1^-/-^,* and *vgll4b^-/-^;vgll4l^-/-^;yap1^-/-^*showed the same number of cells on average, approximately 30% larger than their *yap1^-/-^* siblings and 10% smaller than the *vgll4b^+/+^;vgll4l^+/+^;yap1^+/+^*sibling controls (Fig. 5J-Q, R).

The fact that *vgll4b^-/-^;yap1^-/-^ and vgll4l^-/-^;yap1^-/-^* pLLP are larger than *yap1^-/-^*suggests that Vgll4b and Vgll4l are not “just” competing with Yap1 for Tead binding. These results indicate that Vgll4b and Vgll4l limit the pLLP cell count also independent of Yap1, possibly by actively repressing the expression of genes that promote proliferation ^15, 22^.

To confirm that Vgll4b and Vgll4l have a function independent of Yap1, we overexpressed *vgll4b* and *vgll4l* in the absence of Yap1 to test whether this would cause a further decrease in cell number. Surprisingly, injection of *vgll4b* in *yap1^-/-^*mutant embryos led to a strong, early morphological phenotype, with a very high proportion of crippled embryos compared to the injection in their WT siblings (Fig. 5S-V’). While this strong morphological phenotype did not allow us to quantify pLLP cell number, it clearly showed that Vgll4b is much more potent in the absence of Yap1 protein, further confirming that Vgll4b does not solely function as a competitor of Yap1 to bind to Tead. In the malformed embryos, we noticed that a high number of pLLP largely deviated from their normal migration path, a phenotype that we observed at a low frequency in *yap1* mutant and that we described as “diving” pLLP. While 0,2% of the WT embryos and 16,5% of the *yap1^-/-^*mutant embryos showed “diving” primordia, the proportion was dramatically increased to 54% in *yap1^-/-^* mutants injected with *vgll4b* mRNA (Fig. 5Z). Overexpression of *vgll4l* mRNA in *yap1^-/-^* mutants lead to a similar, but less strong effect with 24% of the embryos showing “diving” primordia (Fig. 5S-T’,W-Y’,Z). These results support the conclusion that both Vgll4b and Vgll4l have Yap1-independent functions.

### 6. Yap1 is active in the pLL placode before migration

Thus far, our data indicate that Yap1 is required to ensure a sufficient number of cells in the pLLP, allowing it to reach the appropriate size. Conversely, the tumor suppressor activity of Vgll4b/4l is required to restrict the number of cells in the pLLP, by acting both as Yap1 competitors and through a Yap1-independent mechanism. We then wanted to determine when Yap1 and Vgll4 are actually active in the lateral line system. For this, we used the *Tg(4xGTIIC:d2EGFP)^mw50^* reporter line in which a destabilized d2GFP is expressed under the control of four TEAD-binding sequences (*GTIIC:d2EGFP*, ^42^). Surprisingly, the reporter activity was very low or undetectable in the migrating pLLP of WT embryos, although it was very high in the surrounding somites (Fig. 6A-A’’). Therefore, we reasoned that Yap1 activity might be higher at 20 hpf, when the pLLP starts to migrate. Indeed, weak and mosaic expression of the reporter could be observed in the pLLP at this stage. (Fig. 6B-B’’). Quantification of the number of *GTIIC:d2EGFP* positive cells confirmed that Yap1 activity was stronger in the pLL before or at the very beginning of migration (Fig. 6F). To confirm that the reporter line was indeed specific for Yap1-Tead transcriptional activity, we used K-975, a small molecule that has recently been described to specifically inhibit YAP1/WWTR1 interaction with TEAD ^43^. The embryos were treated at the 30% epiboly stage and imaged at 20hpf. Treatment with 1µM and 5µM K975 significantly decreased the number of cells expressing the reporter within the pLLP (Fig. 6C-E’’,G). The signal intensity of the TEAD reporter was also significantly reduced in the whole embryo, as well as specifically in the pLLP, after drug treatment (Fig. 6H). The pLLP cell counts in embryos treated with K975 was also significantly reduced (Fig. 6I). We also observed that this effect on cell counts was dose-dependent (Supplementary Fig. 6A-G). Similar results were obtained when treating the embryos at bud stage, past gastrulation (Supplementary Fig. 6H-K). These results indicate that Yap1 is transcriptionally active at the onset of migration and that its activity is likely required before the onset of migration to establish an appropriate number of cells in the placode, and subsequently in the delaminating pLLP.

**Figure 6.**
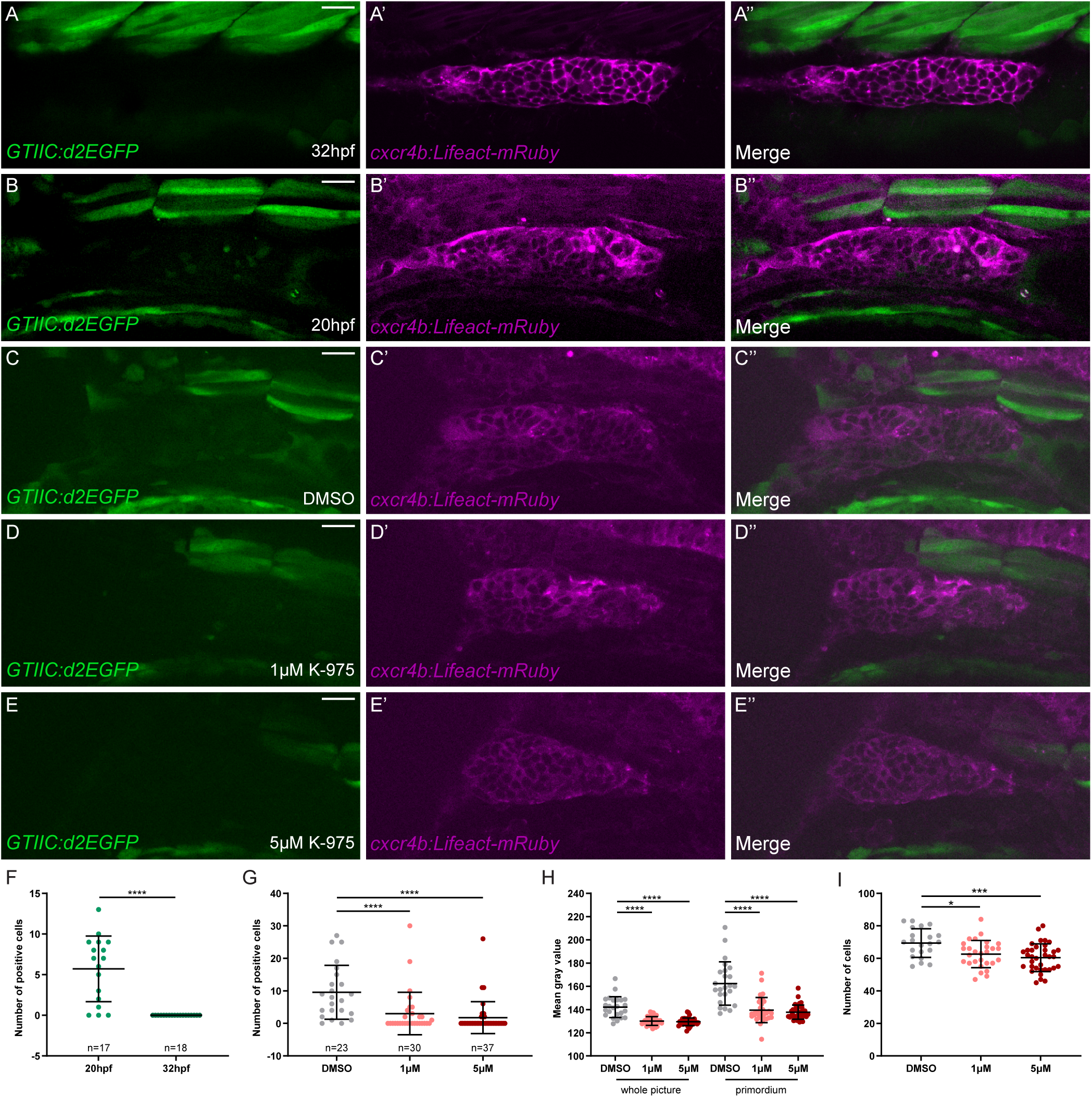
Yap1 transcriptional activity is detected in the pLL placode prior to the onset of migration. (A-B’’) Single-plane confocal images of the pLLP in control embryos at 20hpf (A) and at 32hpf (B) showing *GTIIC:d2EGFP* expression. *cxcr4b:Lifeact-mRuby* is used to visualize the pLLP (A’, B’). Merged images of the two channels are shown in (A’’, B’’). (C-E’’) Single-plane images of confocal Z-stacks of the pLL placode in 20 hpf embryos treated with DMSO (C), 1μM (D) or 5μM K-975 (E) showing *GTIIC:d2EGFP* (C-E), *cxcr4b:Lifeact-mRuby* (C’-E’) and merged channels (C’, E’).. (F, G) Quantification of *GTIIC:d2EGFP* positive cells in the pLLP of 20hpf and 32hpf control embryos (F) and of 20hpf embryos treated with DMSO, 1μM or 5μM K-975 (G). (H) Mean grey value measured either on the whole image or within the primordium on a single plane in the GFP channel in the same embryos for each conditions. (I) Total number of cells in the pLLP in the same embryos for each conditions. Scale bar: 20 μm.

### 7. Vgll4b interferes more than Vgll4l with Yap-Tead transcriptional activity

To further explore the interplay between the Yap1 and Vgll4 proteins, as well as the potential functional differences between the Vgll4b and Vgll4l paralogs, we performed a series of overexpression experiments using the *GTIIC:d2EGFP* reporter line. Overexpression of *yap1* mRNA led to a very strong increase in reporter activity, both across the entire sample (46% increase) and specifically within the pLLP (100% increase) (Fig. 7A-B’’,G-H, Supplementary Fig. 7A-B). In contrast, overexpression of *vgll4b* mRNA resulted in a slight reduction of the GTIIC signal by approximately 10% (Fig. 7C-C’’, G-H, Supplementary Fig. 7A-B). Overexpression of *vgll4l* RNA at the same effective concentration had no noticeable effect on the GTIIC reporter (Fig. 7 D-D’’,G-H, Supplementary Fig. 7 A-B). Notably, co-injection of *vgll4b* with *yap1* led to a significant reduction of the GTIIC signal compared to injection of *yap1* alone (compare Fig. 7E’ to 7B’). Similarly, co-injection of *vgll4l* with *yap1* also reduced the GTIIC signal relative to *yap1* alone, though to a lesser extent than *vgll4b* co-injection (compare Fig. 7F’ to 7B’ and 7E’). Taken together, these findings confirm that Vgll4b and Yap1 compete for binding to Tead, with Vgll4b effectively outcompeting Yap1 and inhibiting its transcriptional activity. Furthermore, these results confirm that Vgll4l functions in a similar manner, albeit with lower potency than Vgll4b.

**Figure 7.**
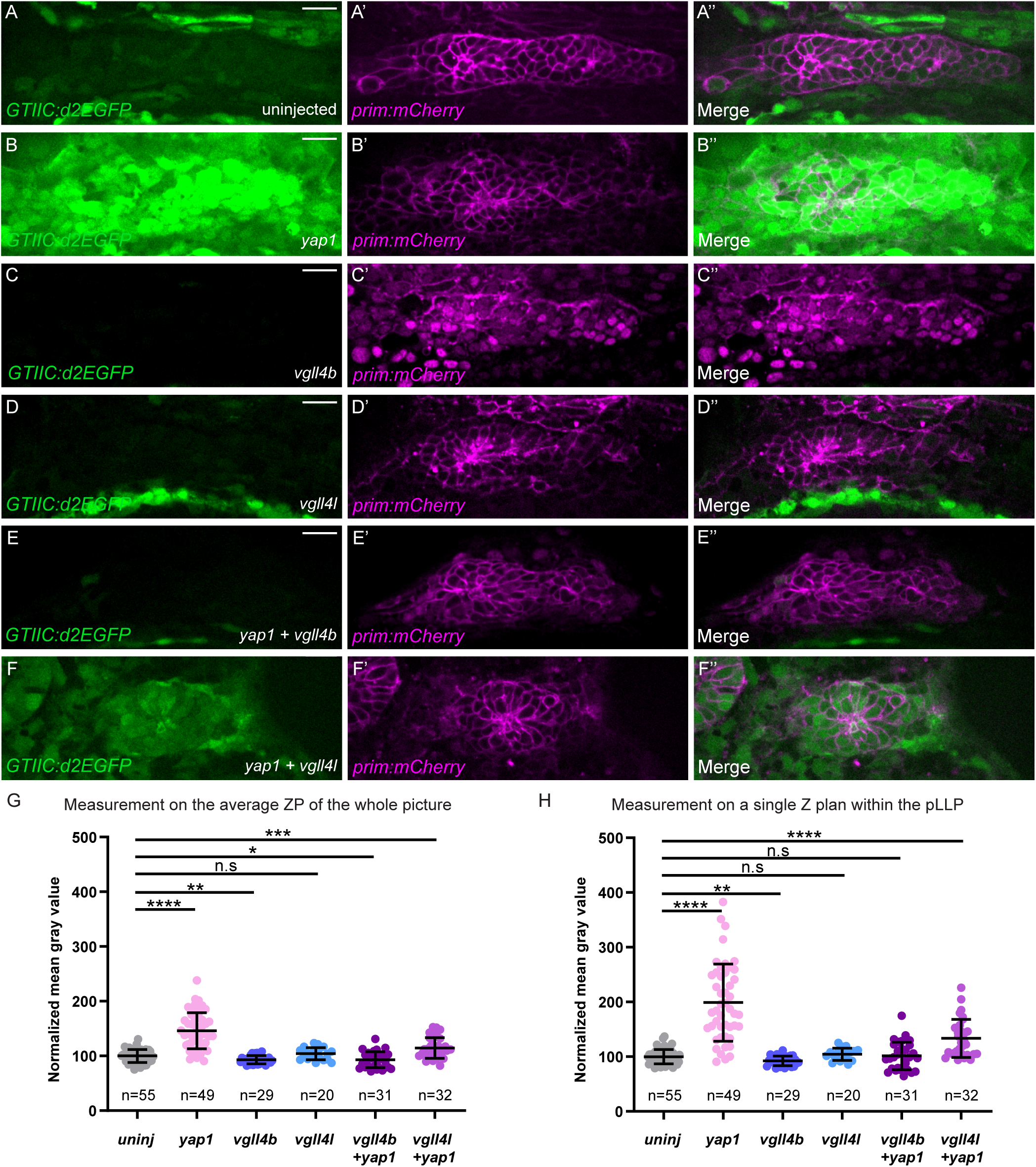
Vgll4b suppresses Yap1-TEAD activity more efficiently than Vgll4l in the pLLP. (A-F’’) Single planes confocal images showing the expression of *GTIIC:d2EGFP* in the pLLP of 20hpf embryos, uninjected (A) or injected with *tBFP-yap1* (B), *tRFP-vgll4b* (C), *vgll4l* (D), *tBFP-yap1* and *tRFP-vgll4b* (E), *tBFP-yap1* and *vgll4l* (F) mRNA. The prim-mCherry transgene is used to visualize the pLLP (A’-F’). Merged images of the two channels are shown in (A’’-F’’). (G, H) Quantification of the mean intensity of *GTIIC:d2EGFP* measured on the average Z-stack Projection of the whole picture (G) or on a single plane within the pLLP in the same embryos (H). Scale bar: 20 μm.

## Discussion

In this study, we reveal that Yap1 growth-promoting activity in the pLL system is regulated not only in the cytoplasm but also in the nucleus by Vgll4 proteins. We propose a model in which Yap1, in association with Tead proteins, ensures that the pLLP contains a sufficient number of cells before migration begins. Conversely, Vgll4b and Vgll4l function partially redundantly to limit the pLLP cell number, with Vgll4b exhibiting a stronger tumor-suppressor activity. Our findings indicate that Vgll4b and Vgll4l counteract Yap1 activity both by competing for Tead binding and through a Yap1-independent mechanism. Overall, this study demonstrates that a precise balance between Yap1 and Vgll4 proteins ensures proper regulation of cell number in the pLLP.

### 1. Yap1 proliferation-promoting activity depends on Tead proteins in the lateral line

During embryonic development, spatio-temporally coordinated cell proliferation is necessary for well-proportioned organ formation and growth. In the pLLP, Wnt/β-catenin signaling is essential for maintaining a population of proliferating progenitors in the leading region during migration ^25, 26^. This is necessary to partially compensate for the loss of cells due to neuromast deposition, ensuring that the pLLP can properly deposit a complete set of lateral line organs ^28, 29^. We have previously shown that the Hippo signalling pathway effector Yap1 is required to ensure that the pLLP contains the appropriate cell number ^30^. Here, we show that in the absence of Yap1 activity, the pLLP is smaller from the onset of migration, suggesting that Yap1 is required prior to migration, while it may also play a role during the migration process. We previously showed that the cell junction-associated protein Amotl2a limits pLLP size, in part by inhibiting the activity of Yap1 ^30^. In the present study, we uncovered a new level of regulation for Yap1 activity in the posterior lateral line. YAP1 is a transcriptional cofactor that depends on DNA-binding proteins to activate gene expression. Although transcription factors of the TEAD family are the main DNA-binding partners, YAP1 can also physically interact with other transcription factors, such as the RUNX2, p73, AP-1, β-CAT ^44–46^. Here, we show that the proliferation-promoting function of Yap1 in the lateral line system is fully Tead-dependent, since (i) the Tead-binding defective *yap1^bns22^* allele shows the same reduction in cell count as the null mutant alleles and (ii) a Tead-binding defective Yap1 was unable to restore the cell number when injected in *yap1* mutants.

### 2. Vgll4b/4l limit cell count in the lateral line system by competing with Yap1

VGLL4 functions primarily by competing with YAP1 to bind to TEAD proteins, physically interacting with TEAD in the same region as YAP1. This competition inhibits the formation of the YAP1-TEAD transcriptional complex in a dose-dependent manner, thereby suppressing the activation of genes involved in cell proliferation and survival ^15, 17, 19, 21, 47, 48^. Our findings align with the established role of VGLL4 in mammals, with Vgll4b and Vgll4l similarly competing with Yap1 for binding to Tead, thereby restricting Yap1 growth-promoting activity in the pLL. First, we show that Vgll4b activity in the pLL depends on the TDU2 motif, which has been shown to mediate the interaction of VGLL4 with TEAD ^15, 21^. Second, the increase in cell number in Vgll4 mutants is largely rescued by the additional loss of Yap1 (in double and triple mutants), indicating an increase in Yap1 activity upon loss of Vgll4 activity. Third, we show that Yap1 is more potent when overexpressed in *vgll4* mutants than in WT, again strongly suggesting that Vgll4 limits Yap1 activity. Altogether, our results indicate that, together, Vgll4b and Vgll4l limit the number of cells and the size of the pLLP by competing with Yap1 to limit its capacity to form an active transcriptional complex with Tead DNA-binding proteins. Which of the four Tead proteins are partners of Yap1, Vgll4b and Vgll4l in the LL system remains to be elucidated.

### 3. Vgll4b/4l also function independently of Yap1

However, our results suggest a more complex role for Vgll4 in regulating cell number in the pLLP. Indeed, if Vgll4b and Vgll4l were to function solely as competitors of Yap1, then their loss- or gain-of-function in *yap1* mutants would have no additional effect over the loss of *yap1* alone. In contrast, we observed that the loss of either *vgll4b* or *vgll4l* alone, as well as their combined loss, can partially rescue the reduced pLLP cell counts observed in *yap1* mutants (Fig. 5J-R). This suggests that Vgll4b and Vgll4l have functions independent of Yap1 in the pLLP. Consistent with this finding, overexpression of *vgll4b* in *yap1* mutants resulted in severe early morphological defects that were not present when *vgll4b* was overexpressed in WT, indicating that Yap1 limits Vgll4b activity. Interestingly, we observed a significant increase in the number of “diving” pLLP upon *vgll4b* overexpression in *yap1* mutants, a phenotype observed at low penetrance in *yap1* mutants (Fig. 5S-Z). This suggests that this phenotype may result from an increase in endogenous Vgll4b activity in the absence of Yap1. Altogether, these results indicate that Vgll4b and Vgll4l do not function solely as Yap1 competitors but also limit cell number in the pLLP in a Yap1-independent manner.

These results are consistent with previous studies showing that VGLL4/Tgi primarily functions as a default repressor in certain contexts, with YAP1/Yki competing with VGLL4 in a dose-dependent manner. For example, Drosophila *tgi* mutants ^19^ and mice lacking VGLL4 function in the liver ^22^ exhibit no overt phenotypes. However, in both cases, the loss of Tgi/VGLL4 fully rescue the Yki/YAP1 loss-of-function, demonstrating that the primary role of Yki/YAP1 in these contexts is to counteract Tgi/VGLL4-mediated default repression ^19, 22^. Similarly, we show that the loss of Vgll4 activity partially rescues the reduction in cell number observed in *yap1* mutant pLLP. Our data support a dual role for Vgll4b/4l in the pLLP, where they function both as default repressors and as Yap1 competitors to regulate cell number and size. Moreover, our results highlight the importance of maintaining a precise balance between Yap1 and Vgll4 binding to Teads, which is likely modulated by factors such as their relative abundance, as well as their binding affinities for Teads.

An alternative, non-exclusive hypothesis is that Vgll4 proteins compete not only with Yap1 but also with another factor, potentially its homolog Wwtr1. Although Wwtr1 alone is not required to control the pLLP size, we cannot rule out the possibility that its function is required in the absence of Yap1, especially since it is expressed in the pLLP during migration (^30^ and data not shown). Yap1 and Wwtr1 function redundantly for posterior body axis morphogenesis and epidermal basal cells in zebrafish ^31, 49^. This could also be the case in the pLLP, although this remains challenging to test due to the severe early morphological defects of the *yap1;wwtr1* double mutant embryos. Wwtr1 could partially account for the Yap1-independent Vgll4 function in the pLLP. In *yap1* mutants, Vgll4b and Vgll4l may still limit Wwtr1 activity, thus accounting for the partial rescue of the reduced pLLP cell count in *yap1* mutant upon additional loss of *vgll4b/4l*.

### 4. Overlap and difference between Vgll4b and Vgll4l functions

The zebrafish genome encodes three Vgll4 paralogs - *vgll4a*, *vgll4b* and *vgll4l*. Previous studies have reported the expression of *vgll4b* and *vgll4l* in the pLL, suggesting their possible roles in regulating cell migration and/or proliferation. In particular *vgll4b* is strongly and continuously expressed throughout migration and in deposited neuromasts ^39^. Our findings support this hypothesis by showing that Vgll4b and Vgll4l are required and function partially redundantly to limit the number of cells in the pLLP. Interestingly, Vgll4b plays a predominant role compared with Vgll4l. On the one hand, Vgll4b alone is necessary and sufficient to limit the number of cells in the pLLP, not Vgll4l alone (Fig. 3G-M). On the other hand, a twofold lower amount of vgll4b mRNA, compared with vgll4l, is sufficient to rescue the number of cells in the pLLP upon total loss of Vgll4 activity (Fig. 4O-U). This functional difference is consistent with sequence homology analysis indicating that, among the three zebrafish paralogs, Vgll4b is much closer to the human VGLL4 protein than Vgll4l (70% versus 32% identity, ^39^). This higher degree of conservation suggests that Vgll4b could also have the highest affinity for Tead and therefore the greatest propensity to compete with Yap1 for binding to Tead. This hypothesis is supported by our findings showing that upon coexpression with Yap1, Vgll4b is more potent than Vgll4l in interfering with Yap1-Tead transcriptional activity measured by the *GTIIC:d2EGFP* reporter (Fig. 7). Further investigations into the specific molecular mechanisms and signaling pathways underlying the distinct functions of Vgll4b and Vgll4l will be crucial in elucidating their precise roles within the Yap1-Tead regulatory axis. Comparative studies examining the binding kinetics, cofactor associations, and target gene regulation of the two Vgll4 isoforms could provide valuable insights.

## Material and Methods

### Zebrafish Husbandry and generation of mutant and transgenic lines

Procedures involving animals were carried out according to the guidelines of the Goethe University of Frankfurt and were approved by the German authorities (Veterinary Department of the Regional Board of Darmstadt). Embryos were staged as previously described ^50^. *yap1^fu47^*, *yap1^fu48^* ^30^, and *yap1^bns22^* ^31^ mutant lines and *Tg(-8.0cldnb:lynEGFP)^zf106^* (*cldnb:gfp*) ^51^, *Tg(4XGTIIC:d2EGFP)^mw50^* ^32^, *Tg(prim:mem-mCherry)* (*prim:mCherry*) ^52^ transgenic lines were characterized previously.

The *vgll4l^fu93^* allele was generated using TALEN targeting exon 2 as previously described in ^30, 53^. The *vgll4b^bns23^*allele was generated using CRISPR-Cas9 with the following sgRNA - 5’- GCAGTCAGCAACCACCGGAC-3’ targeting exon 2 of the *vgll4b* gene. sgRNA DNA oligos were cloned into the pT7-gRNA vector (Addgene plasmid #46759) and *in vitro* transcribed with the MEGAshortscript T7 transcription kit (Ambion). *CAS9* mRNA was obtained by in vitro transcription of linearized *pT3TS-nCas9n* vector (Addgene plasmid #46757) with MEGAscript T3 transcription kit (Ambion). 100 pg of sgRNA and 150 pg of *CAS9* mRNA were co-injected into 1-cell stage AB embryos. The *Tg(cxcr4b(FOS):Lifeact-mRuby)^fu16^* (*cxcr4b:lifeact-mRuby*) was generated using BAC homologous recombination by replacing the second exon of *cxcr4b* in the *cxcr4b*-containing fosmid CH1073-406F3 by *Lifeact-mRuby* followed by zebrafish transgenesis as described in ^54, 55^.

### RNA extraction, cDNA synthesis and RT-PCR

RNA extraction was performed using TRIzol reagent according to manufacturer’s instructions. Embryos at 32hpf were dechorionated and homogenized in 500μL TRIzol with a blunt needle tip syringe (20-40 embryos per tube). cDNA was generated using the iScript^TM^ cDNA Synthesis Kit (Bio-Rad, 1708890) for qPCR or the Invitrogen Superscript III or IV for RT-PCR according to manufacturer’s instructions. The cDNA was diluted 1:2 to obtain a final concentration of 25 ng/μl when used for primer efficiency assays or 1:4 to obtain a final concentration of 12.5 ng/μl when used for comparative gene expression analyses.

Primers were designed using Primer-BLAST and the online OligoAnalyzer tool from Integrated DNA Technologies (IDT) to identify the most optimal primer pairs ^56^. Primer efficiency tests were performed to determine the ability of primers to reliably and accurately bind to the correct sequence ^57^. cDNA diluted for primer efficiency assays was further serially diluted 1:2 to obtain a total of six concentrations (25, 12.5, 6.25, 3.125, 1.5625 and 0.78125 ng/µL). For each primer pair, all concentrations were tested in triplicates. Real-time qPCR was performed using the Bio-Rad iTaq Universal SYBR Green Supermix (1725270) in a CFX Connect Real-Time System (Biorad). Reaction mixes and programs were standardized for all primers which are listed in Table S1. Primer sequences used for the qPCR experiments are listed in Table S1. Fold changes were calculated using the ΔΔCt method ^58^. Expression of the genes of interest were normalized to that of Rpl13.

### Cloning full length and mutated forms of *yap1, vgll4b, vgll4l* and ISH probes

All primers used for cloning are listed in Table S1. Full-length cDNAs encoding *yap1*, *vgll4b* and *vgll4l* and partial cDNA encoding *vgll4a*, *vgll4b* and *vgll4l* were amplified using 30hpf whole embryo cDNA. PCR products were ligated into pCS2-derived vectors (for full-length cDNA) or in pGEM-T (Promega) according to manufacturer’s instructions (for ISH probes). *yapS54A*, *vgll4bΔTDU1* and *vgll4bΔTDU2* were generated by overlapping PCR using the primers listed in Table S1. Capped mRNAs were transcribed with the Sp6 mMessage mMachine kit (Invitrogen, AM1340).

### Anti-Yap1 Antibody Production

To generate the antibody against zebrafish Yap1, the fragment of Yap1 encoding the first 230 amino acids was cloned into the expression vector pGST-MCS-6XHIS. The resulting pGST-Yap1 (1-230)-6XHIS was then produced in large quantities (BIOSS Toolbox Core Facility, Freiburg, Germany) and shipped to Eurogentec (Belgium) to immunize two rabbits pre-selected for their low pre-immune serum background. After the immunization program, different serum samples were received and directly tested by immunostaining on WT and *yap1* mutant embryos.

### Whole-Mount Immuno-fluorescence and Whole Mount In Situ Hybridization (WISH)

For whole-mount immunostaining, embryos were fixed overnight at 4°C or room temperature for 2h in 4% PFA in 1X PBS (pH=7.4). After fixation, embryos were washed with PBS-T (PBS + 0.1% tween-20) for 5X 5 minutes. Fixed embryos were used directly for the experiments without upgrading them in methanol. The rest of the procedure was performed according to standard procedures ^59^. In this study, rabbit anti-Yap1 (1:200, VL7801), mouse anti-GFP (1:500, Takara Bio Europe/France, JL8), and rabbit anti-pH3 (1:200, Santa Cruz Biotechnology, Dallas, TX, USA) were used as primary antibodies. Anti-rabbit and anti-mouse Alexa conjugated antibodies were used as secondary antibodies (1:500, Molecular Probes, Eugene, OR, USA).

Whole-mount *in situ* hybridization was performed according to standard procedures ^59^. The *vgll4a*, *vgll4b* and *vgll4l* probes were used at 1:200 dilution. To visualize the pLLP after WISH, an immunostaining with the mouse anti-GFP antibody was performed as explained above.

### Image acquisition

#### ISH images

Imaging of ISH embryos was performed on an inverted microscope (Ti-S) using a 4x objective to acquire whole embryos and a 20x objective for close-ups of the pLLP.

#### Imaging

Imaging of live and fixed embryos was performed on a Nikon W1 Spinning Disc Microscope with the following objectives: 10× air objective (NA 0.45, WD 4 mm), 20× (NA 0.95, WD 0.95 mm), and 40× (NA 1.15, WD 0.60 mm) water objectives. The embryos were mounted as described in ^60^, with the following modifications: no second layer of 0.5% LMPA was added on top of the 0.3% LMPA. The image recording software was NIS-Elements v4.

#### Image processing and quantification of cell numbers in the pLLP

Unless indicated otherwise, all cell counts were performed when the primordium had reached the middle of the yolk extension in control and mutant embryos (approximatively 32hpf). Cell counts at an earlier developmental stage were performed on embryos between 18 and 22ss. All images were processed using Fiji software.

For fixed samples and injections of full-length and mutated forms of *yap1* in the *yap* mutants (Figure 2, Supplementary Figure 2) the number of cells in the pLLP was counted manually using the Cell Counter plugin.

For all other live acquisitions, automated cell counting was performed using the GFP signal of the *cldnb:gfp* transgenic embryos. Single cells of the primordium were segmented in 3D using a homemade Fiji plugin based on MorphoLibJ and 3D manager built-in plugins. First, the plugin registers the pLLP in X and Y, and cropping in Z. This is performed by generating a maximum Z-projection, which is then blurred. The image is segmented in 2D using a minimum threshold and rotated by the angle formed by the difference between the long axis of the ellipsoid and the horizontal axis. After the rotation, the image is cropped. Once the pLLP is registered, single-cell segmentation starts on the Z-stack image using the MorphoLibJ plugin after a step of 3D Gaussian blur. Filter and clearing is then performed to remove segments below or above a certain volume threshold and clear blank slices in Z. Finally, single-cell measurements are obtained using the 3D Manager plugin and saved in the output files. Manual checking of the quality of 3D segmentation was systematically performed. Inaccurate segmentations (under or over segmentation) were removed from the analysis.

R v3.6.1 was used to compile the segmented data. A homemade R script allowed for the removal of individual cells from the dataset, which were likely not part of the pLLP (wrongly segmented skin cells). The detection of these outliers was based on volume and surface/volume ratio. The cell count and total volume of the primordium were extracted from these datasets.

#### Neuromast deposition

Neuromast deposition pattern analysis was performed using a custom Fiji macro script that segments individual cell clusters and the pLLP. The images were segmented based on optimized filter parameters derived from trial and error. After segmentation, the macro allows for manual correction. The ROIs and their corresponding positions were saved in output files. R v3.6.1 was used to compile cell cluster data. Using, a homemade R script, the positions of the neuromasts were normalized to the initial detected cell cluster (ganglion), and the number of neuromasts deposited was calculated. Additionally, the number of neuromasts deposited at the end of migration was counted manually on independent datasets.

#### Rosette Detector

The method used for rosette detection is based on a convolutional neuronal network (CNN) ^61^ and was modified from the “rosette detector” algorithm previously used in the laboratory and described in ^62, 63^. The algorithm was updated with a state-of-the-art CNN using Caffe as a backend ^61^ and re-trained with manually labeled primordia. The script was launched in Visual Studio Code (V1.53.2). Using maximum Z-projection images as an input, the algorithm detects structures similar to the rosettes and assigns them a score indicative of similarity to the wild-type training data. The number of detections and weighted (sum of detection scores) detections were saved in a .csv output file. R v3.6.1 was used to calculate the ratio of weighted detections over the number of detections to reflect the “rosettiness” of a pLLP.

#### Aspect ratio measurement

Aspect ratio measurements were performed by manually encircling the pLLP on maximum Z-projection images using the freehand selection tool in Fiji. Shape descriptors were selected in the Set Measurements menu which yielded measures of circularity, aspect ratio (AR), roundness, and solidity.

For all analyses, processed data were extracted from R as .csv files and imported into GraphPad Prism for plotting.

### Statistical analysis

GraphPad Prism was used for statistical analysis, and samples were subjected to the two-tailed unpaired t-test (also called Mann-Whitney test) to determine the *P-*values. *P*-values are shown in the figure panels as ****p<0.0001 and ^ns^p>0.05. In the graph, data are represented as mean±SD.

### Single Embryo Genotyping

Embryos were retrieved from mounting stamps; as described in ^60^. Single fixed/live embryos were boiled at 95^°^C in 50mM NaOH (Stock 1M in ddH_2_O) for 1h and then neutralized by adding 1/10^th^ of the volume of 1 M Tris-Cl (pH 8.2). After NaOH neutralization, 2μl of this crude preparation was used for PCR genotyping ^30, 53^. Genotyping of *yap^fu47^* and *yap^fu48^* and *taz^fu55^*were performed as described previously ^30, 53^. Genotyping of *vgll4b^bns23^*, *vgll4l^fu93^* and *yap^bns22^*were performed using primers listed in Table S1. The *vgll4l* and *yap^bns22^*PCR products did not require further digestion and were loaded directly onto 4% and 3% agarose gels, respectively. The *vgll4b* PCR product was digested with PstI overnight before loading onto a 2% gel.

## Acknowledgments

We thank Dr. Holger Knaut for the prim-mCherry transgenic line and the BIOSS Toolbox Core Facility (Freiburg, Germany) for help with the production of GST-Yap1 protein. We are grateful to the animal caretaker team for excellent fish care, to S. Link, M. Kamprad, A. Niedztwezki and M. Heyde for technical assistance, and to our bachelor and master students M. Porsiel, N. Sheth and T. Urosevic for their help. We thank L. Desruelles and A. Mertens for critical reading of the manuscript. This work was supported by the German Research Foundation (LE2681 in SPP1782 to C.D and V.L. and Walter Benjamin Program to A.L.), the Humboldt Research Fellowship for postdoctoral researchers, the Fondation Bettencourt Schueller and the Add-on Fellowships for Interdisciplinary Life Science (Joachim Herz Stiftung) [to A.L.].

## Author Contributions

A.L., C.D., V.D. designed and performed most of the experiments, analyzed the data, and wrote the manuscript. P.A.K, C.H., C.S. O.M. and S.P. performed and analyzed some experiments. D.K. and T.F. helped with the analysis. J.K.K.L in the lab of D.Y.R.S generated the *yap1^bns22^* and the *vgll4b^bns23^* mutant lines. V.L. coordinated the project, helped to design experiments, supervised the work and wrote the manuscript together with the authors.

## Conflict of Interest

The authors declare they have no conflict of interest.

---

**Supplementary Figure 1.**
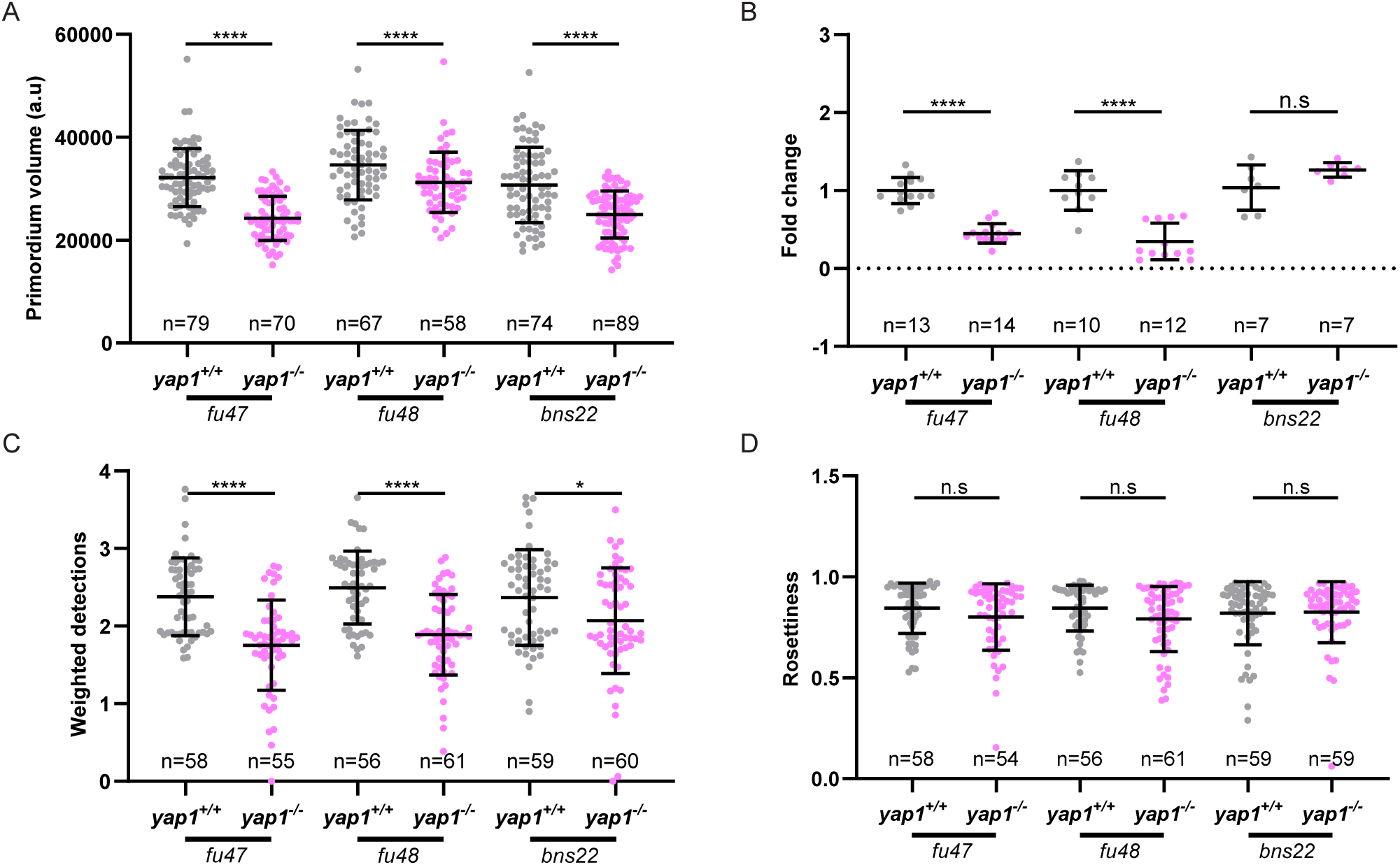
Tead-binding is essential for Yap1-driven proliferation in the pLLP. (A) Relative *yap1* transcript levels measured by qPCR in *yap1^+/+^*and *yap1^-/-^* siblings carrying the *fu47*, *fu48* or *bns22* alleles at 32hpf. (B-D) Quantifications of total volume (B), rosette weighted detections (C) and rosettiness index (D) of the pLLP in in *yap1^+/+^* and *yap1^-/-^* siblings carrying the *fu47*, *fu48* or *bns22* alleles at 32hpf.

**Supplementary Figure 2.**
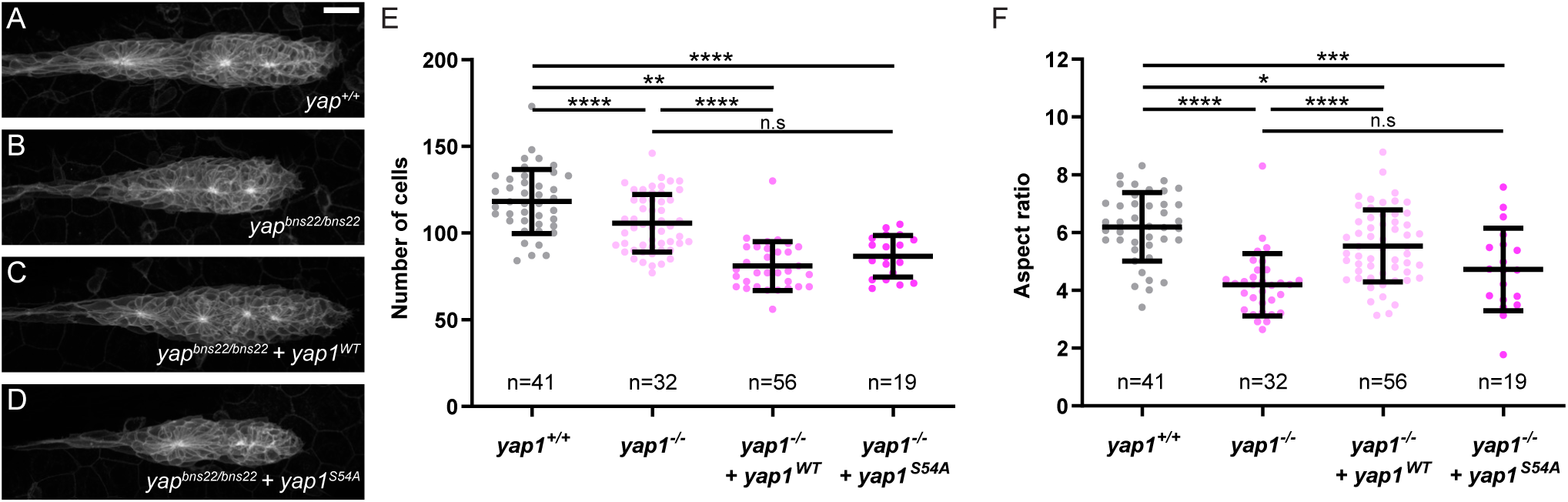
A Tead-Binding-Deficient Yap1 mutant fails to rescue *yap1* loss-of-function phenotype. (A-D) MIP of confocal Z-stacks showing the pLLP in uninjected *yap1^+/+^* (A), uninjected *yap1^bns22/bns22^*embryos (B), *yap1^bns22/bns22^* embryos injected with *yap1*-WT mRNA (C) and *yap1^bns22/bns22^* embryos injected with *yap1-S54A* mRNA (D). (E-F) Quantification of pLLP cell number (E) and aspect ratio (F) in each condition. Scale bar: 20 μm.

**Supplementary Figure 3.**
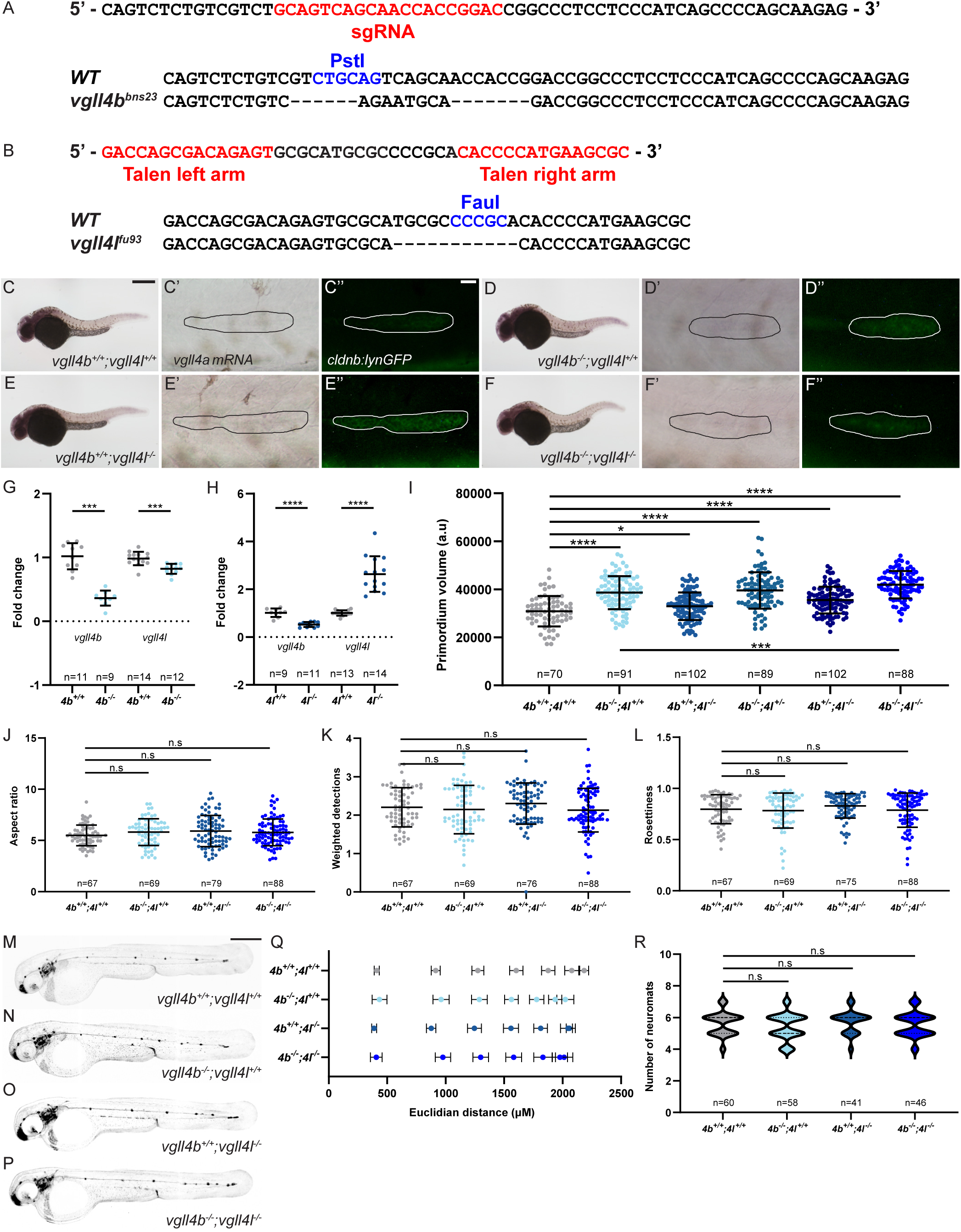
Vgll4b and Vgll4l are required to limit the number of cells in the pLLP. (A) CRISPR target sequence within the *vgll4b* exon2. (B) TALEN target sequence within the *vgll4l* exon2. (C-F’’) Overview images of 32hpf embryos stained with a *vgll4a* ISH probe in *vgll4b^+/+^;vgll4l^+/+^*(C), *vgll4b^-/-^;vgll4l^+/+^* (D), *vgll4b^+/+^;vgll4l^-/-^*(E), and *vgll4b^-/-^;vgll4l^-/-^* (F) embryos. Images of the pLLP at a higher magnification, stained with *vgll4a* (C’-F’) ISH probe and an anti-GFP antibody (C’’-F’’) in the pLLP in the indicated genotypes. (G-H) *vgll4b* or *vgll4l* transcript levels measured by qPCR in *vgll4b^-/-^* (G) and *vgll4l^-/-^*(H) mutants at 32hpf. (I-L) Quantifications of the total volume (I), aspect ratio (J), weighted detections (K), and rosettiness (L) of the pLLP in the indicated genotypes. (M-P) Overview images of 48hpf *vgll4b^+/+^;vgll4l^+/+^* (M), *vgll4b^-/-^;vgll4l^+/+^* (N), *vgll4b^+/+^;vgll4l^-/-^* (O), and *vgll4b^-/-^;vgll4l^-/-^* (P) embryos. (Q-R) Quantifications of neuromast deposition pattern (Q) and total number of neuromasts (R) for each indicated genotype. Scale bar: 400 μm (C-F, M-P), 20 μm (C’-F’’).

**Supplementary Figure 4.**
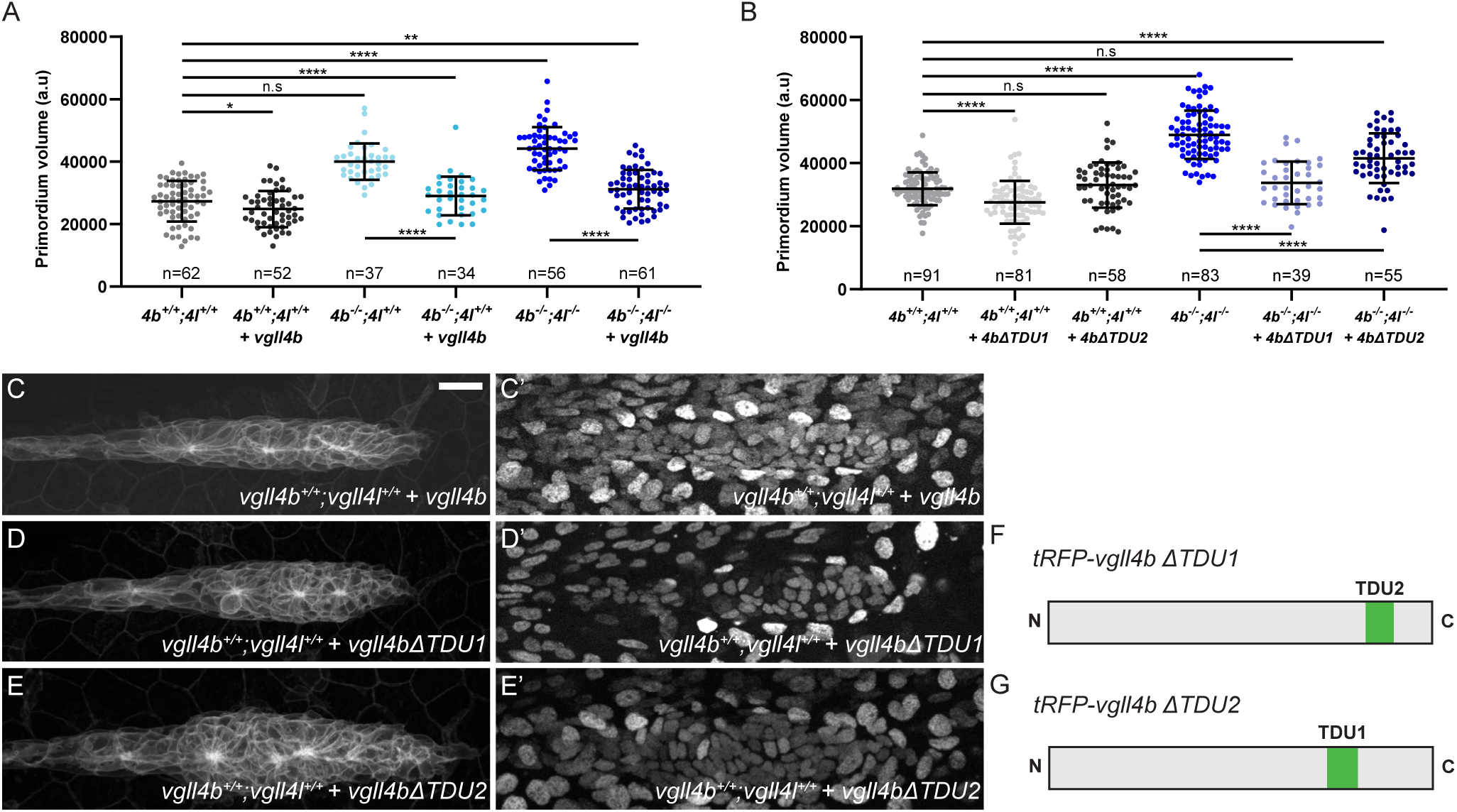
Vgll4b and Vgll4l are sufficient to rescue the loss of Vgll4 activity in the pLLP, and Vgll4b TDU2 is required for this function. (A-B) Quantification of pLLP volume in the indicated genotypes. (C-E’) MIP of confocal Z-stacks showing the pLLP (C-E) and the localization of *tRFP-vgll4b* (C’), *tRFP-vgll4bΔTDU1* (D’) and *tRFP- vgll4bΔTDU2* (E’) injected into *vgll4b^+/+^;vgll4l^+/+^* embryos. (F-G) Schematic representations of the truncated Vgll4b forms lacking TDU1 (F) or TDU2 (G). Scale bar: 20 μm.

**Supplementary Figure 6.**
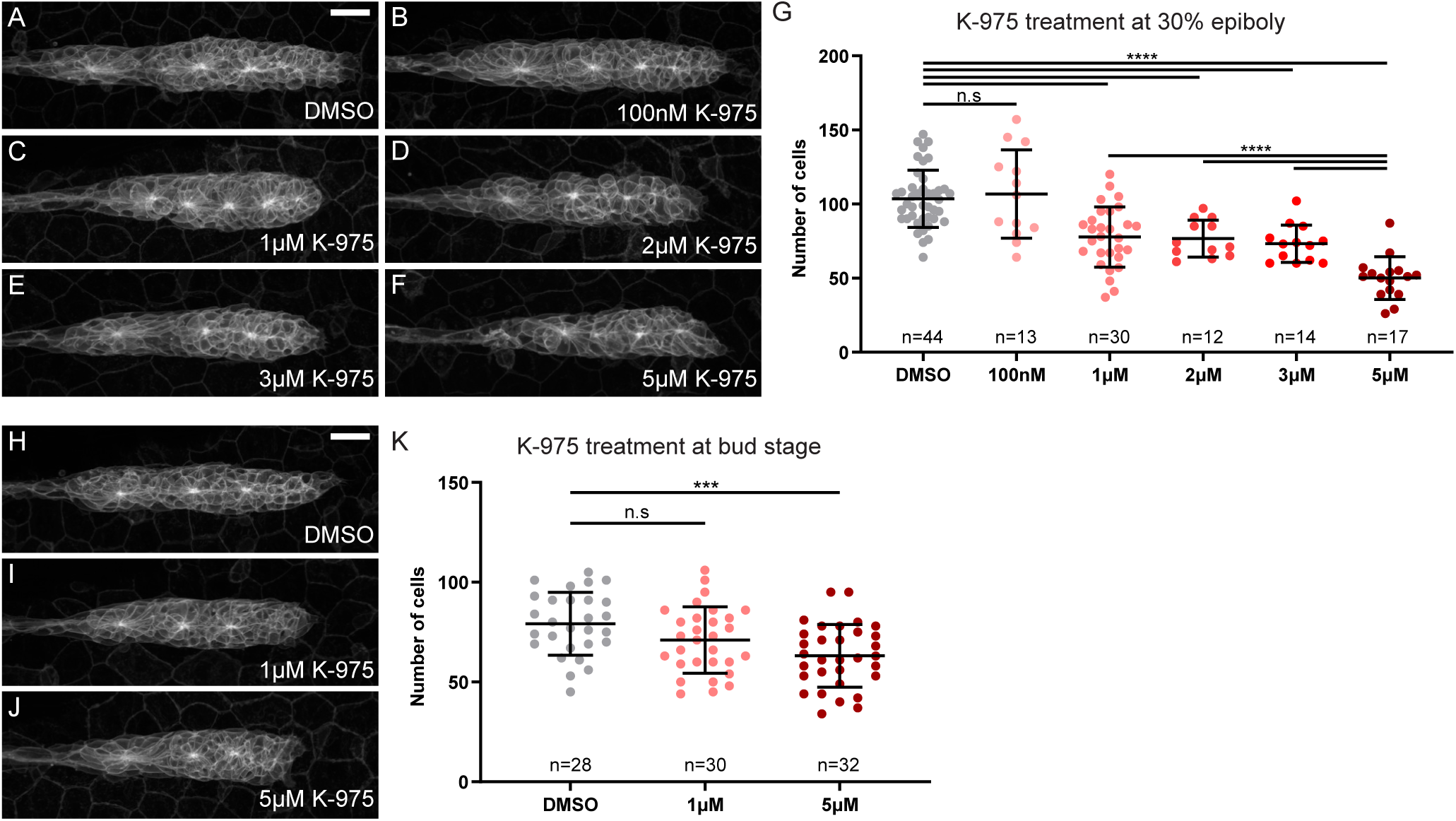
Yap1 transcriptional activity is detected in the pLL placode prior to the onset of migration. (A-F) MIP of confocal Z-stacks showing the pLLP in 32hpf embryos treated at 30% epiboly with DMSO (A) or increasing concentrations of K-975: 100nM (B), 1μM (C), 2μM (D), 3μM (E) and 5μM (F). (G) Quantification of pLLP cell number at 32hpf in the indicated groups. (H-J) MIP of confocal Z-stacks showing the pLLP of 32hpf embryos treated at bud stage with DMSO (H) or 1μM (I) and 5μM (J) K-975. (K) Quantification of pLLP cell number at 32hpf in the indicated groups. Scale bar: 20 μm.

**Supplementary Figure 7.**
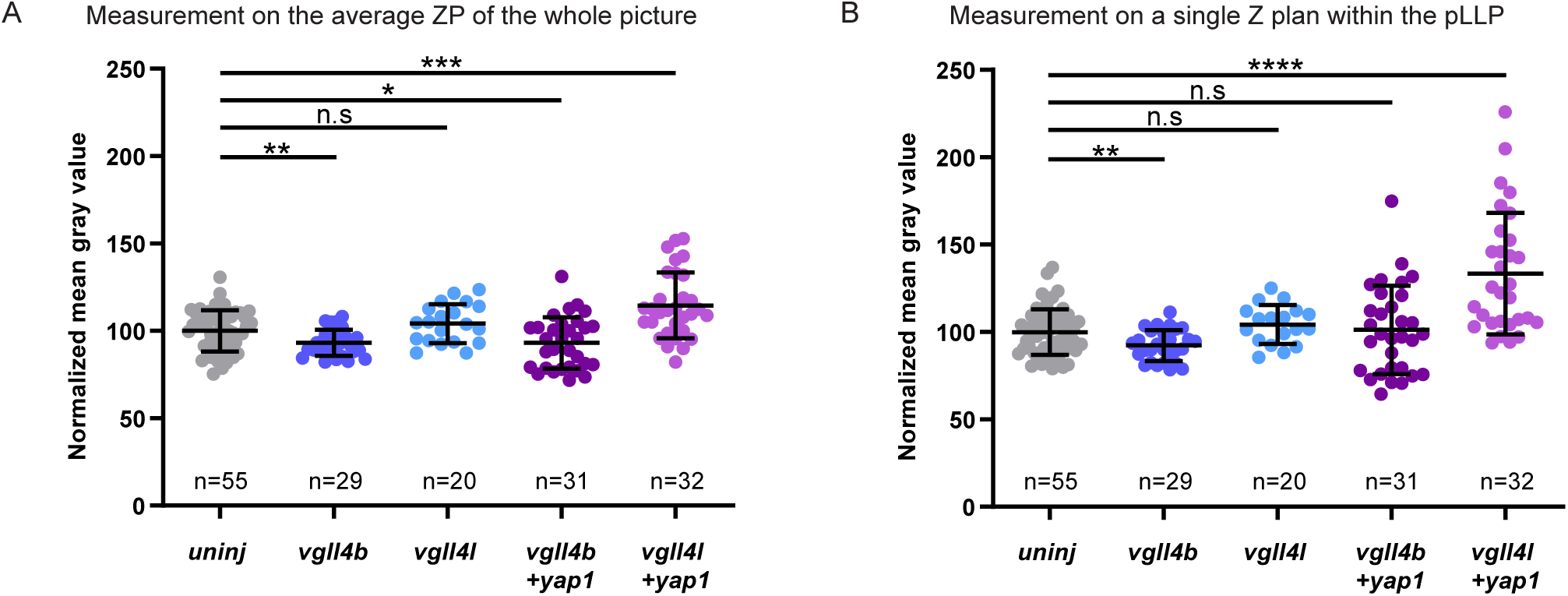
Vgll4b suppresses Yap1-TEAD activity more efficiently than Vgll4l in the pLLP. (A, B) Quantification of normalized *GTIIC:d2EGFP* mean intensity measured on the average Z-Projection of the whole image (A) or within the primordium on a single plane in the same embryos (B).

**Table S1.**
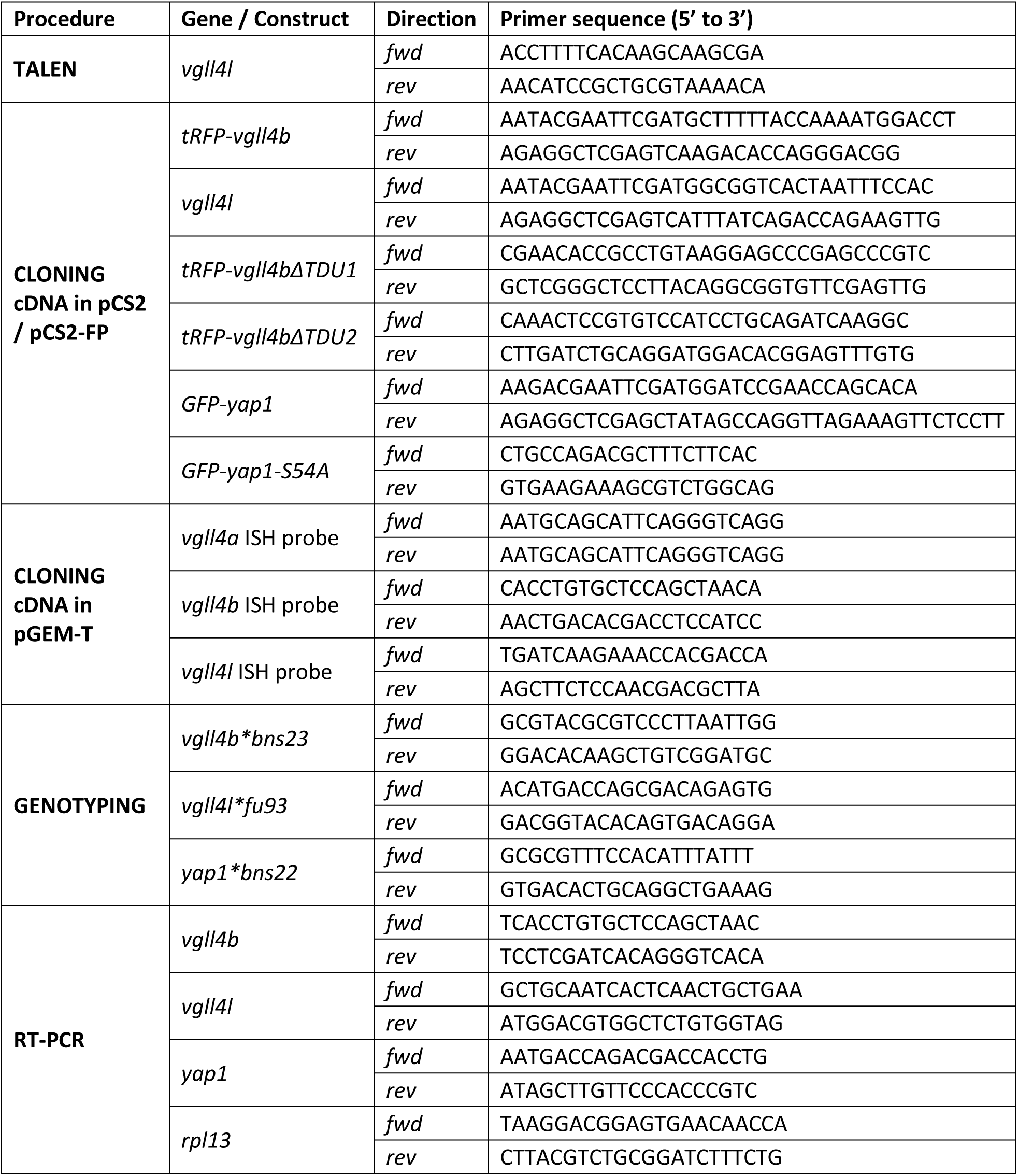

## Notes

### Competing Interest Statement

The authors have declared no competing interest.

